# Cell-type Plasticity Supports Behavioral Adaptations at the Water-to-Land Interface

**DOI:** 10.64898/2026.07.25.740661

**Authors:** Andrew MM Matheson, Jamie Woych, Therese G Zinga, Nat Spollen, Maxime Policarpo, Giacomo Gattoni, Gabriel J Graham, Luke T Geiger, Alonso Ortega-Gurrola, Eliza CB Jaeger, Walter Salzburger, Maria Antonietta Tosches

**Affiliations:** Department of Biological Sciences, Columbia University, New York, New York, USA; Zoological Institute, Department of Environmental Sciences, University of Basel, Basel, Switzerland; Department of Neuroscience, Columbia University, New York, New York, USA

## Abstract

Animals inhabiting aquatic or terrestrial habitats experience different constraints on their physiology and locomotion, and are exposed to fundamentally different sensory environments. Across evolutionary timescales, most species have adapted to live exclusively either in water or on land. Newts are among the vertebrates that defy this rule and split their adult lives between freshwater ponds and terrestrial habitats. In these amphibians, transitions across environments cause remarkable phenotypic plasticity in their body morphology. But whether and how the nervous system and behavior also adapt to these environmental changes remains poorly explored. Here, we establish the Iberian ribbed newt *Pleurodeles waltl* as a new model to study the neurobiology of environmental plasticity in a vertebrate. We first show that experimental transitions between aquatic and terrestrial laboratory settings recapitulate morphological changes observed in the wild. Furthermore, aquatic and terrestrial newts display plasticity in sensory and motor behaviors, including differences in walking gait and odor responsiveness. In the olfactory system, the transition from water to land involves a profound remodeling of the nasal epithelium, including reversible transcriptomic changes in secretory and support cells, and an increase of neurogenesis. Together, our findings reveal how plasticity of specific cell types in the nervous system supports behavioral adaptations across environments. More broadly, this work establishes newts as a model to study the functional constraints and convergent adaptations that may have shaped the evolution of vertebrate nervous systems in water and on land.

## Introduction

Animals are well adapted to the environments that they live in, but some animals inhabit very different environments over their lifetime (Stevens 2016, Westneat et al., 2018). Many species respond to environmental change with a change in phenotype, through a process known as phenotypic plasticity. Classic examples include the coat coloration changes observed seasonally in many birds and in arctic mammals, morphological differences in spadefoot toad larvae, and developmental plasticity of the pectoral girdle in bichir fish reared on land (Zimova et al. 2018, Denver et al., 1998, Standen et al., 2014). How phenotypic plasticity changes the body and improves fitness has been explored in several systems (West-Eberhard, 2003, Sommer 2020), but relatively little is known about phenotypic plasticity of the nervous system and the underlying cellular and molecular mechanisms.

Among vertebrates, dramatic cases of phenotypic plasticity are found in amphibious fishes and newts, animals that inhabit both aquatic and terrestrial environments (Ashley-Ross et al., 2013, Wright et al., 2016). Moving between water and land is one of the most drastic environmental changes an animal can experience, as it requires adjusting metabolism, nitrogen excretion, osmoregulation, and locomotion (Matheson 2025, Escoriza and Hassine 2019, Zamora-Camacho 2023, Kristin and Gvozdik 2014, Walters and Greenwald 1977, Degani and Meerson 2024, Brown et al 1984). Transitions between these two environments also impose distinct demands on sensory processing. Light, sound, and odors all propagate differently through water and air (Steele et al. 2023, MacIver and Finlay 2022, MacIver et al, 2017, Mollo et al., 2014, 2017, Ladich and Winkler 2017). Odor plumes, in particular, have different spatiotemporal statistics that depend on the fluid dynamics of the medium through which they propagate (Connor et al., 2018, Moore and Crimaldi 2004, Waldrop et al., 2018). In addition, chemicals can act as odorants only if they can reach the olfactory epithelium in the nose, and their ability to do so depends on the medium. A chemical may be volatile, soluble, both, or neither, meaning the chemical cue emanating from the same source could have two different signatures at the nose in water and on land (Mollo et al., 2014, 2017).

Life on Earth started in the ocean, and transitions to land were associated with convergent changes in animal genomes (Wei et al., 2026) and major innovations in body plan, physiology, and the nervous system (Vermeij 2017). As a result, most vertebrate species are specialized for either aquatic or terrestrial environments. Studying phenotypic plasticity in lineages that have independently adapted to the water-to-land interface (Ashley-Ross et al., 2013, Wright et al., 2016) can reveal the constraints and selective pressures that have shaped one of the most consequential ecological and evolutionary transitions in vertebrate history.

To determine whether and how the nervous system adapts to environments with extremely different physical properties, we focused on the Iberian ribbed newt, *Pleurodeles waltl*, a species from the Iberian peninsula and North Africa. Newts are a subfamily of salamanders (called *Pleurodelinae*, Fig. 1A, Rancilhac et al., 2021) characterized by a semiaquatic lifestyle. Like most amphibians, their life cycle involves an aquatic larval stage followed by metamorphosis. However, unlike most other salamanders, post-metamorphic newts divide their lives between aquatic and terrestrial habitats (Fig. 1B). For many species of newts, transitions between water and land occur when the ephemeral pools they live in dry up, forcing animals to repeatedly alternate between aquatic and terrestrial lifestyles. Field studies have revealed that these environmental transitions correlate with phenotypic changes, such as the dramatic remodeling of the dorsal crest in modern European newts (Wiens et al., 2011, Zamora-Camacho 2023). Although environmentally induced changes in the newt olfactory system were first documented nearly a century ago (Matthes 1927, Schulze 1934), the plasticity of the nervous system and behavior in these animals has remained largely unexplored.

**Figure 1:**
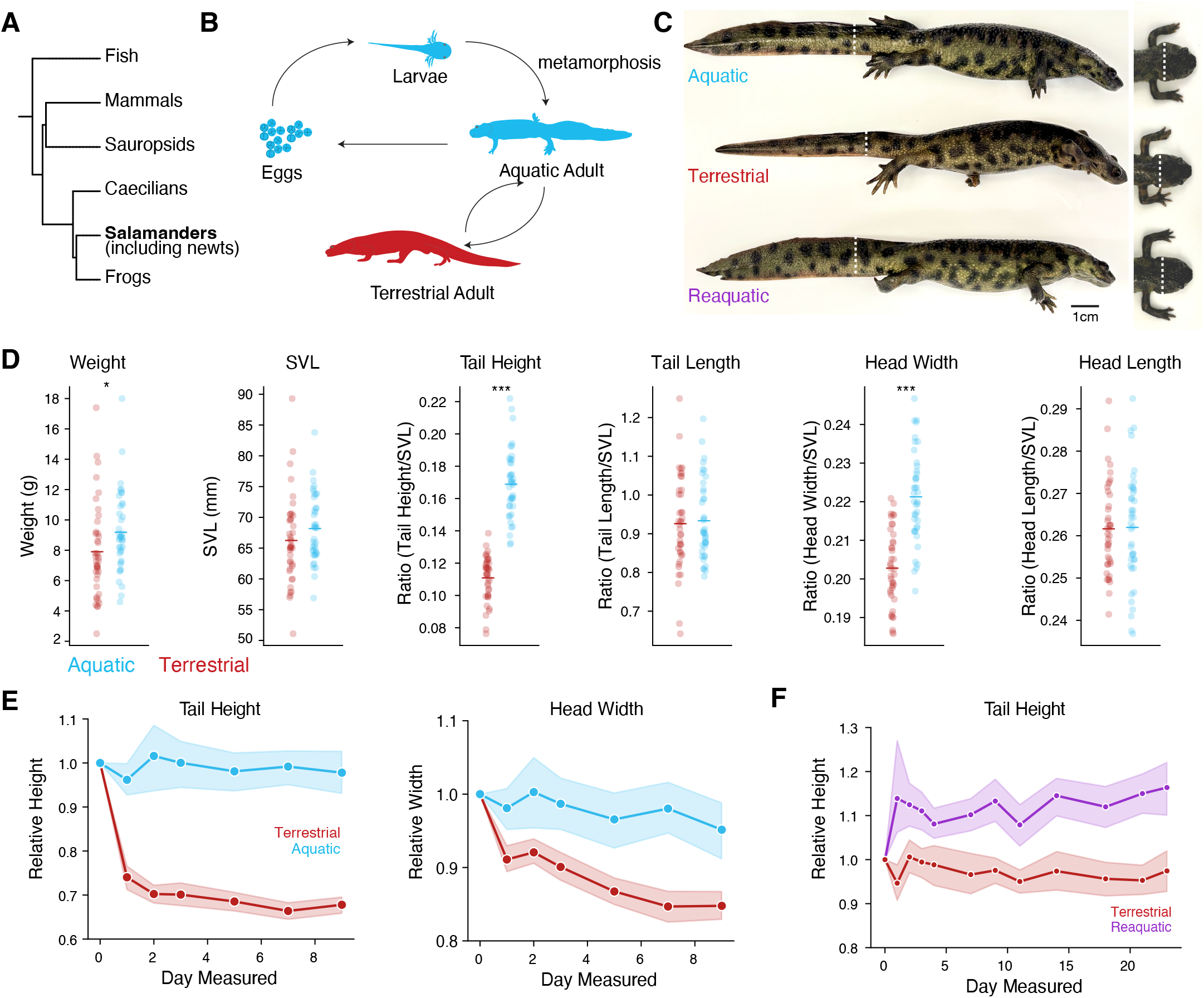
Transitions between aquatic and terrestrial environments trigger rapid and reversible morphological changes in the newt *Pleurodeles*. A) Simplified phylogenetic tree of vertebrates. B) Lifecycle of *Pleurodeles waltl*. Blue: aquatic phase. Red: terrestrial phase C) Photos of *Pleurodeles* post-metamorphic siblings habituated to terrestrial conditions, aquatic conditions, and reaquatic conditions showing tail and head morphology, dashed white line indicates tail height and head width measurement locations. D) Measurements of weight, SVL (snout-vent length: distance from the nose to the bottom of the cloaca), tail height, tail length, head width, and head length across the *Pleurodeles* colony. Height and width normalized to SVL. Tail heights and head widths are significantly different between terrestrial and aquatic conditions (p=1.16e-21,and p=9.14e-10, age-dependent linear model (see Methods), while tail length, head length, and SVL do not differ significantly (adjusted p=5.84e-01, adjusted p=7.51e-01, adjusted p=6.96e-02, age-dependent linear mode). Weight had a significant age dependent difference (adjusted p=2.86e-02, age-dependent linear model). Terrestrial animals: average age at measurement 29 months, transferred on land at 13 months of age on average and kept in terrestrial conditions for 15 months on average before measurement. Aquatic animals: average age at measurement 22 months; N=45 aquatic and N=46 terrestrial in total. E) Time course of tail height and head width reduction, normalized to the measurements on the first day, during the first 9 days of life on land (N=22) and in aquatic controls (N=10). Line: mean; shaded region: 95% confidence interval. F) Time course of tail height change after returning terrestrial animals to water (“reaquatic”, purple, N=10) and in terrestrial controls (N=10). All P-values are corrected for multiple comparisons with the Bejamini-Hochberg FDR procedure.

Here we show that transitions between aquatic and terrestrial habitats in adult newts cause plasticity across levels, from gene expression to behavior. Plastic changes span multiple body systems, and are reversible at each environmental switch. Using a naturalistic olfactory behavior paradigm, we find that newts habituated to water or land respond differently to the odor of their food. Consistent with this, the nasal epithelium undergoes a complete remodelling after the water-to-land transition, with secretory and support cells changing their transcriptomes, and progenitors and stem cells upregulating neurogenesis. Our findings establish *Pleurodeles* as a vertebrate model for understanding the neural bases of behavioral adaptations to environments with different physical properties.

## Results

### *Pleurodeles* rapidly change their body morphology between environments

Since the 1950s, *Pleurodeles waltl* has been maintained in the laboratory using aquatic husbandry protocols (Gallien, 1952, Beetschen 1995). To induce environmental plasticity in the laboratory, we established a “terrestrial” housing paradigm, where post-metamorphic animals (>8 months old) are placed in a humid environment but have no ability to submerge themselves. We found that animals can live and grow for years (up to 4.2 years to date) in these conditions. A broad morphometric analysis of aquatic and terrestrial animals in our colony (11-22 months on land) revealed that terrestrial animals have reduced tail heights and head widths, while other measurements (tail, head, and snout-to-vent lengths) were not significantly different between the two cohorts; terrestrial animals also showed an age-dependent difference in overall bodyweight (Fig 1C-D). These observations demonstrate that morphological differences are induced by the two different husbandry conditions.

To determine the timescale of morphological change, we then measured head widths and tail heights of animals actively undergoing the transition from water to land (Fig 1E). Tail fin reduction was particularly rapid, with >25% of tail height being lost in one day. Finally, we asked whether these morphological changes are reversible once terrestrial animals are returned to water (here called the “reaquatic” cohort). Measurements of the tail fin showed an increase of tail height after return to water, with a ∼10% increase by day 9 (Fig. 1F). These results show that morphological differences induced by our laboratory paradigm recapitulate those described in aquatic and terrestrial newts in the wild (Zamora-Camacho 2023, Degani and Meerson 2024). In addition, we establish the time course of these changes, and show that they are reversible.

### Plasticity of motor behavior

Next, we asked whether plastic changes extended beyond the body and also impacted behavior. Newts, by virtue of their semiaquatic lifestyle, maintain two distinct motor programs throughout their lives: they swim by bending axial muscles when in water, and walk with their limbs when placed on a substrate. This is why salamanders have traditionally been used to study how central pattern generators in the spinal cord control swimming and walking (Delvolvé et al., 1997, Ijspeert et al., 2007, Chevallier et al., 2008, Ryczko et al., 2020). However, it is unknown whether newts adapted to water or land walk in the same way, given that the tail, which affects balance and coordination (Schwaner et al., 2021, Fish et al., 2021), is remodeled across environments.

We placed *Pleurodeles*, habituated to terrestrial or aquatic environments, into a long, narrow enclosure on land and recorded their locomotion from above for 20 minutes. Next, we tracked 21 points along their body using the deep learning pose estimation package Lightning Pose (Biderman et al. 2024) (Fig 2A). These data were then analyzed with Keypoint-MoSeq, an algorithm that clusters body positional data into discrete behavioral modules called “syllables” (Weinreb et al., 2024), and produces data-driven statistics on animal locomotion. This approach identified a variety of behavioral syllables that we could categorize as walking, hunches, bends, head extensions, rearing, or stopping (Fig. 2B,C, Supplemental Fig. 1A). We found that many walking syllables were used at significantly different frequencies across conditions (Fig 2B). To determine if these changes were reversible after an extended terrestrial phase, we examined behavioral syllables in a group of reaquatic animals, which were returned to water for 30 days after spending more than one year on land. The syllable frequencies in reaquatic animals were more similar to aquatic animals than to terrestrial animals (Fig. 2B).

**Figure 2:**
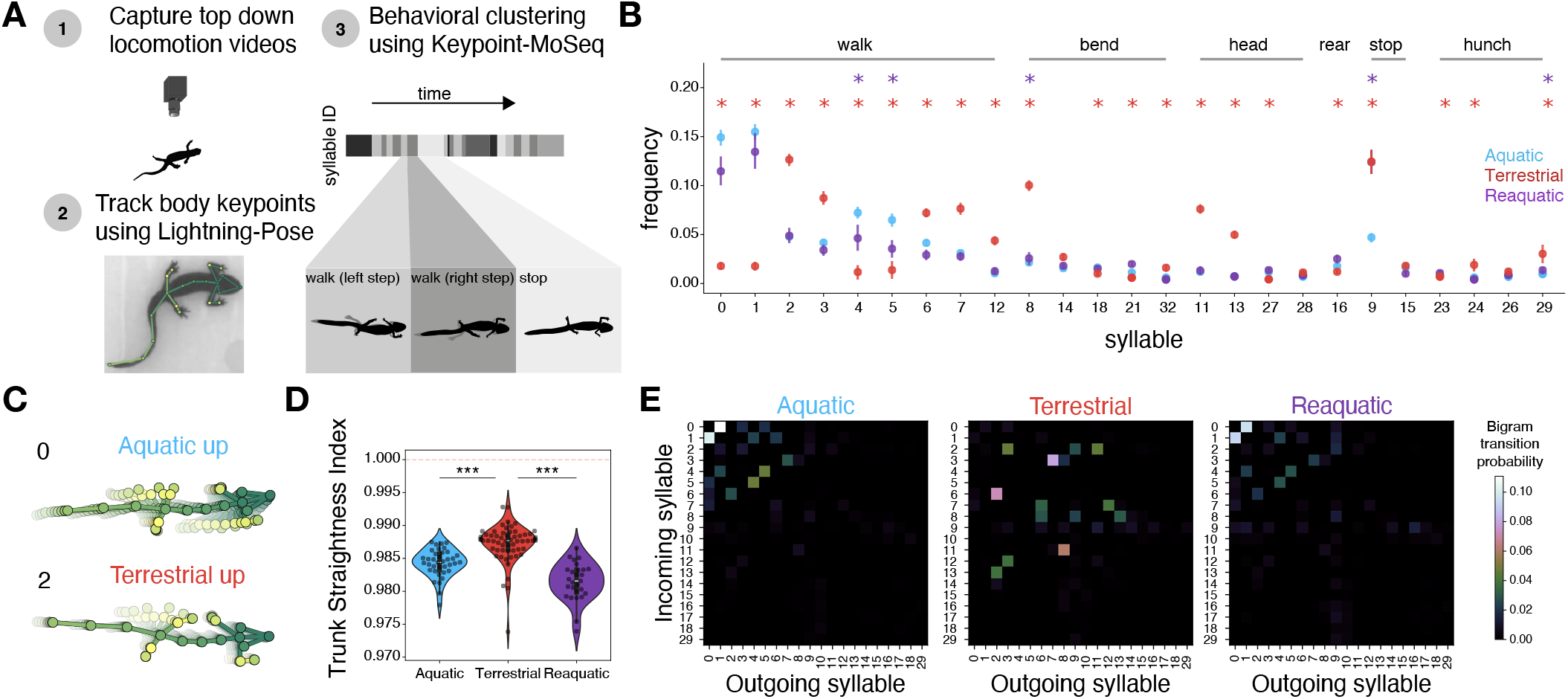
Locomotor gait changes after habituation to aquatic or terrestrial environments. A)Schematic of behavioral video capture and analysis, depicting the 21 keypoints used for tracking overlaid on an individual *Pleurodeles* frame. B) Frequency of syllable use (mean ± SE) in terrestrial (red, N=23, 70 trials), aquatic (blue, N=21, 37 trials), and reaquatic animals (purple, N=15, 28 trials), syllables sorted by behavioral category. Significance stars indicate syllables come from significantly different distributions (Kruskal-wallis test p<0.05). Red: terrestrial vs aquatic, purple: reaquatic vs aquatic. C) Visualization of example walking syllables that are used differently in terrestrial and aquatic animals. D) Violin plots of the straightness index (straight line distance between the neck and base of tail keypoints divided by the total spine length) during walking bouts in aquatic (N=21, 37 trials), terrestrial (N=21, 58 trials), and reaquatic animals (N=15, 27 trials) after filtering for sufficiently consistent walking . Bars: Interquartile range; white line: mean. The trunk of terrestrial animals is significantly straighter than in aquatic and reaquatic animals (p=1.90e-06, p=6.21e-05, linear mixed effects model). E) Transition bigrams depicting probability of transitioning from incoming syllables (rows) to outgoing syllables (columns) in terrestrial, aquatic and reaquatic, displaying only syllables with >1% frequency overall in the data. All P-values corrected for multiple comparisons using the Benjamini-Hochberg FDR procedure.

Overall, these differences in syllable frequency indicated that aquatic animals bend their trunks, move their heads, and sway their tails more than terrestrial animals, which instead maintain a forward gait and keep their trunks and tails straight. To quantify this more directly, we segmented all walking bouts from the data and computed the average trunk straightness index, defined as the ratio of the distance between the base of the neck and the base of the tail, and the total trunk length (sum of all four keypoint segments between the neck and the base of the tail). Consistent with our observations, terrestrial newts had significantly higher trunk straightness indices compared to aquatic or reaquatic newts, indicating a different usage of axial muscles across environments (Fig. 2D).

Finally, we asked whether locomotion in aquatic and terrestrial animals differs not only in relative syllable usage, but also in the patterns of syllable sequences. By examining the transition probabilities between syllables, we identified patterns of syllable sequences that were characteristic of terrestrial or aquatic locomotion; the transition probabilities between syllables for aquatic and reaquatic animals were more similar to each other than to terrestrial animals (Fig 2E). These results indicate that locomotion undergoes plastic changes when animals transition from water to land, and that these changes are reversible upon return to an aquatic environment.

### Environment-dependent modulation of olfactory behavior

After observing the plasticity of motor systems, we wondered whether environmental transitions may similarly impact sensory systems. Several considerations prompted us to focus on olfaction. In amphibians, olfaction plays a major role in food search, navigation, and mating behaviors (Vyatkin and Shakhparonov, 2024, Malacarne and Vellano 1982), as reflected by the prominence of their nasal cavity and olfactory processing brain areas (Woych et al., 2022). Given that odor identity and propagation dynamics differ drastically across media (Mollo et al., 2014, 2017, Steele et al. 2023), we also reasoned that plasticity of the olfactory system may be adaptive. We thus designed a series of behavioral experiments to test responses to airborne odorants and compare newts housed in water to those adapted to a terrestrial environment for at least 6 months.

Newts adjust their breathing frequency by expanding and contracting their buccal cavity, pulling air in and out of their nose. The presence of odors causes an increase of the frequency of their buccopharyngeal oscillations, enabling active sampling of the local odor space (Joly and Cuillère, 1983, Jørgenson 2000). We thus asked whether exposure to an ethologically-relevant food odor (midge larvae, also known as bloodworms) on land modulated buccal breathing frequency in terrestrial and aquatic animals. To measure buccal breathing, newts were filmed from four directions simultaneously, while allowed to explore a terrestrial arena; buccal movements were then scored using BORIS (Fraird and Gamba, 2016) during exposure to odorless airflow (wind) or food odor. Only terrestrial newts increased their breathing rate between the wind and odor phases, indicating that aquatic newts did not respond to the food odor in these experimental conditions (Fig. 3A).

**Figure 3:**
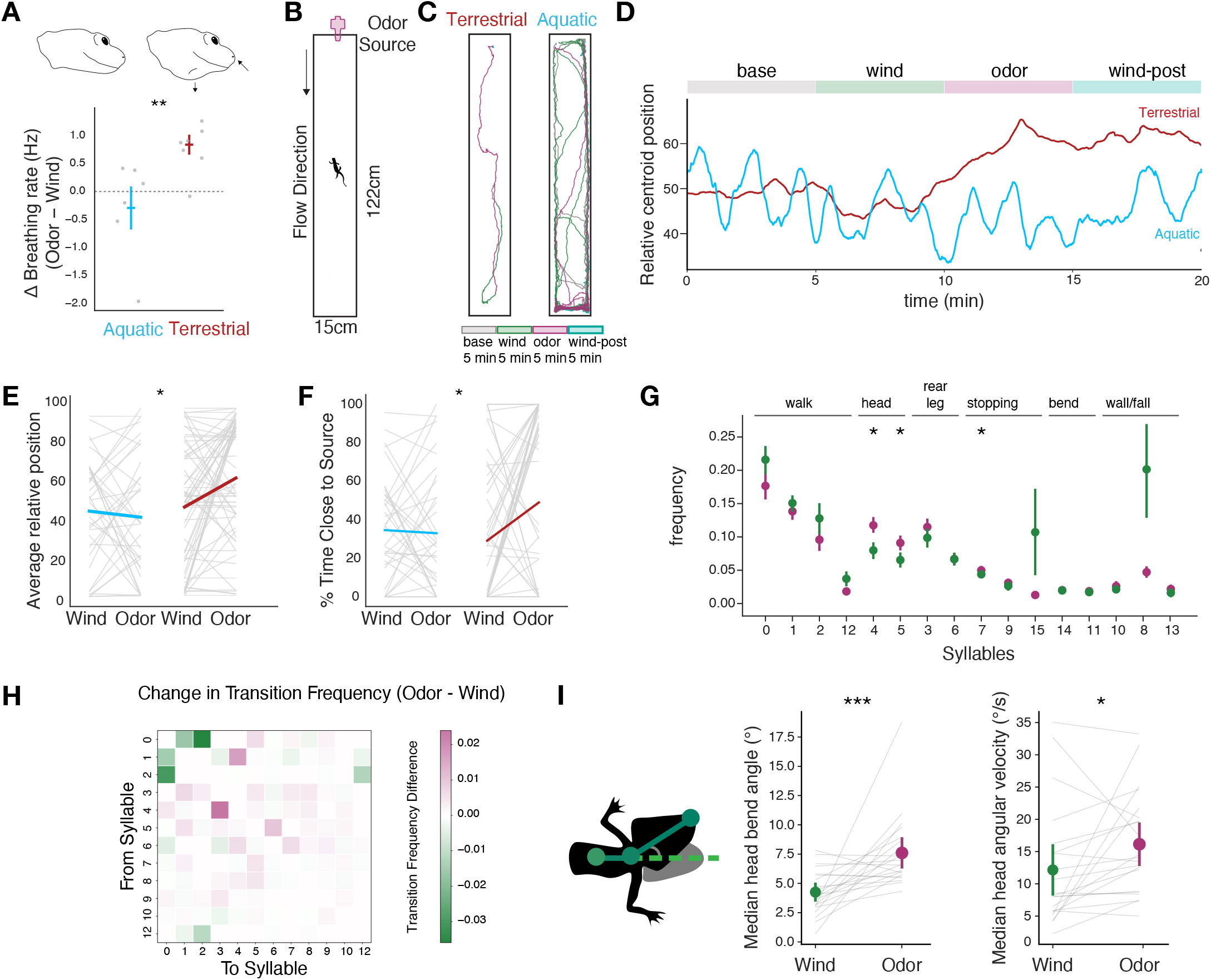
Behavioral responses of terrestrial and aquatic newts to food odor presented on land. A) (top) Newts draw air in their nasal cavities by expanding the floor of their buccal cavity. (bottom) Modulation of breathing rate in aquatic (N=6) and terrestrial animals (N=7), measured by comparing the end of the wind phase and start of the odor phase. Difference in breathing rate change is significantly higher in terrestrial animals (p=4.66e-3, Mann-Whitney U test). Grey dots represent individual animal differences while horizontal bars represent means with error bars of ± 1SE. B) Schematic of terrestrial wind tunnel arena with dimensions and flow direction of the odor plume. C) Traces color coded by stimulus for example terrestrial and aquatic newts. D) Average relative position of aquatic (N=21,37 trials) and terrestrial newts (N=23, 70 trials) along the long axis) of the arena 50 indicates the center, higher positions are closer to the odor source. E) Average relative arena position for aquatic (blue, N=21, 37 trials) and terrestrial newts (red, N=23, 70 trials), measured during the last 2.5 minutes of odor and wind phases of the trial. On average, terrestrial animals move closer to the odor source during the odor phase, while aquatic animals do not change their average position (adjusted p=1.1e-2, linear mixed effects model). F) Time spent close to the odor source (top 1/4) by aquatic and terrestrial newts during the last 2.5 minutes of the wind and odor phases. Terrestrial animals spend more time closer to the odor source after odor presentation compared to aquatic ones (p=1.8e-2, linear mixed effects model). G) Frequency of syllable use (mean ±SE) in terrestrial animals during wind and odor periods (N=16 animals, n=23 trials). Significance stars denote syllables with significantly altered usage during odor relative to wind (adjusted p<0.05, general linear mixed effects model). H) Transition bigram depicting the difference in syllable transition probabilities during the wind and odor phases of the trial (in terrestrial animals), with positive values (magenta) indicating syllable transitions common during the odor period. I) (left) schematic showing head angle measurement. (middle) increase of head bend angle of terrestrial newts during the odor phase of the trial compared to the wind period (right) increase of head angular velocity in terrestrial newts during the odor phase compared to the wind period (p= 2.27e-05, p=3.40e-2, respectively, linear mixed effects model). All p-values corrected for multiple comparisons using the Benjamini-Hochberg FDR procedure.

To test whether olfactory-driven behaviors on land require adaptation to the terrestrial environment, we developed a food odor plume tracking assay for *Pleurodeles*. Newts were placed in a linear track (122cm x 15cm, Fig 3B) where olfactory stimuli were delivered from a nozzle at one end of the arena. After a baseline period, animals were exposed to clean airflow, then food odor, and finally clean airflow again (5 minutes each, for a total trial length of 20 minutes). Overall, we found that terrestrial *Pleurodeles* moved upwind towards the source of the odor plume at the onset of the odor presentation, and spent more time in the area closest to the odor source (Fig. 3C-F). In contrast, aquatic animals did not move towards the odor source during the odor period, suggesting that they could not detect the food odor in this terrestrial assay (Fig. 3C-F). Next, we asked whether terrestrial animals exhibited signatures of active search behavior during the odor presentation phase. We thus trained a new Keypoint-MoSeq model on the data from terrestrial newts that moved upwind during the odor period (Supplemental Fig. 2A). We found that during the odor phase there was an increased frequency of syllables related to stopping and head motion, and that walking syllables were more likely to transition to non-walking syllables, suggesting an increase in odor search behavior (Fig 3G-H). In line with these observations, we also found that the angle between head and trunk, and the head angular velocity, increased in the odor phase compared to the wind phase of the trial, when newts were facing the odor source (Fig 3I). Overall, this suggests that terrestrial newts use a typical olfactory search algorithm, similar to the one described in mammals (Baker et al., 2018): they go upwind towards the odor source and then periodically stop to actively sample the odor plume by moving the head left and right.

Together, these findings align with classical work by Matthes (Matthes, 1927) who found that aquatic *Triturus* newts cannot detect food in terrestrial environments. Our behavioral results show that aquatic-habituated newts do not react to olfactory stimuli when placed on land, suggesting that plastic adaptations of the olfactory system are needed to detect odors in a terrestrial environment.

### Environmental change causes gene-expression changes in the nasal epithelia

Next, we asked whether cellular and molecular changes in the nervous system underlie the environmental plasticity of olfactory behaviors we observed. To investigate this, we began at the sensory periphery, the nasal epithelia. The newt nasal cavities are shaped as simple, unfolded sacs extending from the nostrils rostrally to the choanas (openings on the roof of the mouth) caudally. The nasal epithelium comprises an olfactory epithelium organized in rostrocaudally oriented stripes separated by wedges of respiratory epithelium, in addition to a secretory epithelium adjacent to the vomeronasal organ (Matthes 1927, Schultze 1934, Nakada et al., 2014, Weiss et al., 2021, Reiss and Eisthen 2008, Rozanski and Zuwala, 2019). These regions could be easily identified in *Pleurodeles* (Fig 4A-D, Supplementary Fig. 3, Finlay et al., 2022, Deprez et al., 2020, Durante et al., 2020, Brann et al., 2020). Histological analyses of the nasal epithelia from aquatic and terrestrial animals revealed differences between conditions, including an increase in the number of Bowman’s glands in terrestrial newts (Supplementary Figure 4).

**Figure 4:**
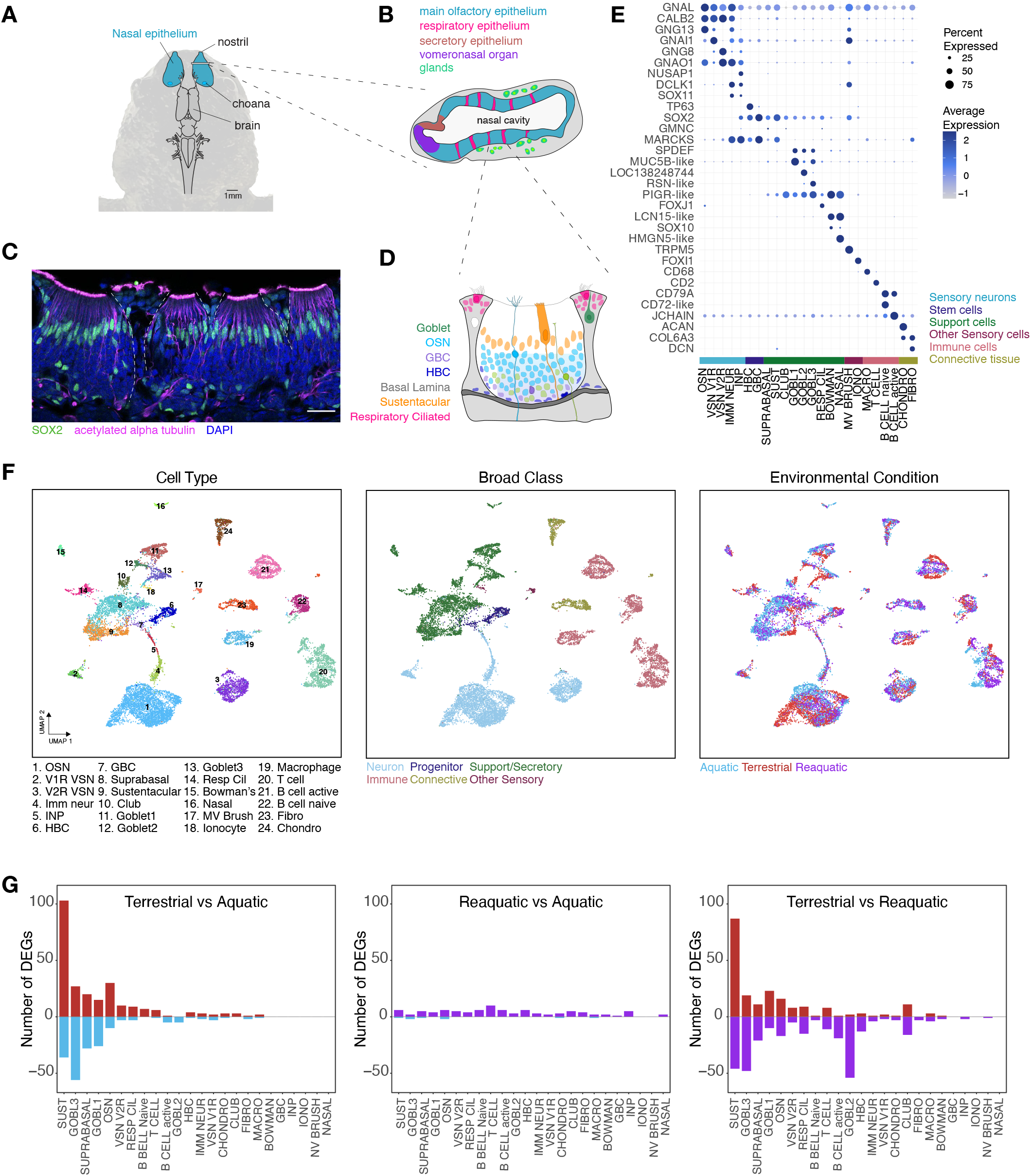
Transcriptomic profiling of the nasal epithelium across environments. A) Dorsal view on the *Pleurodeles* head depicting the nasal epithelia (blue) and the brain. Drawn to scale, scale bar 1mm. B) Schematic of a coronal section of the nasal cavity showing its internal organization, including the main olfactory epithelium (blue), the respiratory epithelium (magenta), the vomeronasal epithelium (purple), the secretory epithelium (brown) and the secretory glands (green). C) Immunostaining for SOX2 (green) to mark the nuclei of sustentacular cells (superficial) and of stem cells (deeper in the epithelium), and acetylated alpha tubulin (magenta) to mark sensory neuron processes and cilia. Blue: DAPI staining (nuclei). Scale bar: 200um. Dashed white lines depict the border of the olfactory epithelium stripes and respiratory epithelium wedges D) Schematic diagram showing one stripe of olfactory epithelium flanked by wedges of respiratory epithelium, with major cell types depicted in different colors. Stem and progenitor cells (HBCs, navy, and GBCs, purple) are found at the base of the epithelium, above the basal lamina. Immature sensory neurons (green) are more abundant at the base of the olfactory epithelium. Mature sensory neurons (blue) extend an apical process to the surface of the epithelium, which branches out into multiple cilia. Sustentacular cells (orange) have a microvillar apical surface facing the nasal cavity, anchor on the basal lamina, and have superficial elongated nuclei. Respiratory multiciliated cells (magenta) in the respiratory epithelium wedges line the nasal cavity. Goblet cells and Bowman glands (not shown) secrete components of the mucus (dark green and green). E) Dotplot showing marker genes for each cell type annotated in the dataset, with cells clustered by broad class (side annotation bar). Dot size represents percent of cells expressing the gene, color represents average expression. F) Uniform manifold approximation projection (UMAP) plots of the nasal epithelia scRNA-seq merged dataset color coded for cell type (left), with numeric annotations below, broad class (middle), and environmental condition (right). Each dot represents the transcriptome of a single cell, n=14,739. G) Bar plots showing the number of differentially expressed genes for each cell type and across conditions, positive values indicate the gene is upregulated in the first condition, negative values indicate the gene is upregulated in the second condition (MAST with logFCthreshold=1.5, minimum change in percentage of cells expressing the gene of 20%).

We hypothesized that three potential changes in the nasal epithelia may modulate the ability to detect odorants in water and air. First, olfactory and vomeronasal sensory neurons (OSNs and VSNs) may express different sets of chemoreceptors, specialized in detecting waterborne or airborne chemicals in the two environments. Second, sensory neurons may simply increase in number in terrestrial newts to enhance odor sensitivity on land, where odors diffuse more rapidly from the source. Third, odor detection might be modulated by changes in the extracellular environment of sensory neurons, which affects odor diffusion, binding to olfactory receptors (ORs), and sensory transduction (Pelosi et al., 1996, Bryche et al., 2021).

To probe the existence of cellular and molecular changes in the nasal epithelium upon environmental transitions, we performed single-cell RNA sequencing (scRNA-seq) in steady-state aquatic, terrestrial, and reaquatic *Pleurodeles*. After quality filtering, we obtained 15,813 single-cell transcriptomes, and then integrated data from all animals to conservatively annotate cell types (Supplemental Fig. 3A). The integrated dataset included 64 clusters of neuronal and non-neuronal cells (Supplemental Fig. 3A), which were annotated using well-established marker genes from human, mouse, and fish datasets (see Fig. 4E, Supplemental Fig. 3 and 5, Finlay et al. 2022, Brann et al., 2020, Deprez et al 2019, Durante et al., 2020, Takaoka et al., 2025, Chen et al., 2025).

The nasal epithelium included all major cell types described in the olfactory and respiratory epithelia of other vertebrate species: secretory cell types that produce mucus, respiratory ciliated cells, OSNs and VSNs, support cells called sustentacular cells (Fig. 4B,C), and horizontal and globose basal stem and progenitor cells (HBCs and GBCs, respectively) (Fig. 4B,D, Supplemental Fig. 5). Embeddings of the non-integrated data suggested that the aquatic clusters were more similar to the reaquatic condition than to terrestrial clusters (Fig. 4F). To systematically identify the effects of environmental changes on nasal epithelium cell types, we computed the number of differentially expressed genes (DEGs) across conditions for each cluster using MAST, a conservative statistical framework (Finak et al., 2015). This analysis revealed over one hundred DEGs between aquatic (or reaquatic) and terrestrial animals, but a minimal number of DEGs between aquatic and reaquatic animals. Notably, sustentacular cells were the cell type with the largest number of DEGs across conditions, followed by goblet cells, a type of secretory cell (Fig. 4G).

To gain insights on the functional changes associated to these transcriptomic shifts, we performed gene expression enrichment analysis using the AUCell algorithm for each cell type against the Hallmark50 dataset from the Molecular Signatures Database (MSigDB) (Liberzon et al., 2015), which groups genes by functional categories. Terrestrialization produced a marked upregulation of interferon alpha and gamma response genes across most cell types, indicating epithelial stress (Fig. 5A). Notably, sustentacular and goblet cells displayed cell-type specific expression enrichment for additional gene sets (Fig. 5A). In contrast, OSNs and VSNs did not show any unique cell-type specific transcriptomic change. These observations point to sustentacular and goblet cells as major players in the adaptation of the nasal epithelium between water and land.

Sustentacular cells form the barrier between the environment and the olfactory epithelium, and are marked by the expression of SOX2 (Fig 4C,D). Besides interferon response genes, these cells displayed increased expression of reactive oxygen species pathway, oxidative phosphorylation, and xenobiotic metabolism genes in the terrestrial condition (Fig 5A,B). Notably, terrestrial sustentacular cells expressed high levels of the inducible nitric oxide synthase NOS2 (Fig. 5B,C), suggesting modulation of innate immunity and mucociliary clearance processes (Bayarri et al., 2021). These transcriptomic changes were reversible upon return to water (Fig 5B), indicating that sustentacular cells switch between two distinct molecular states depending on the environment.

**Figure 5:**
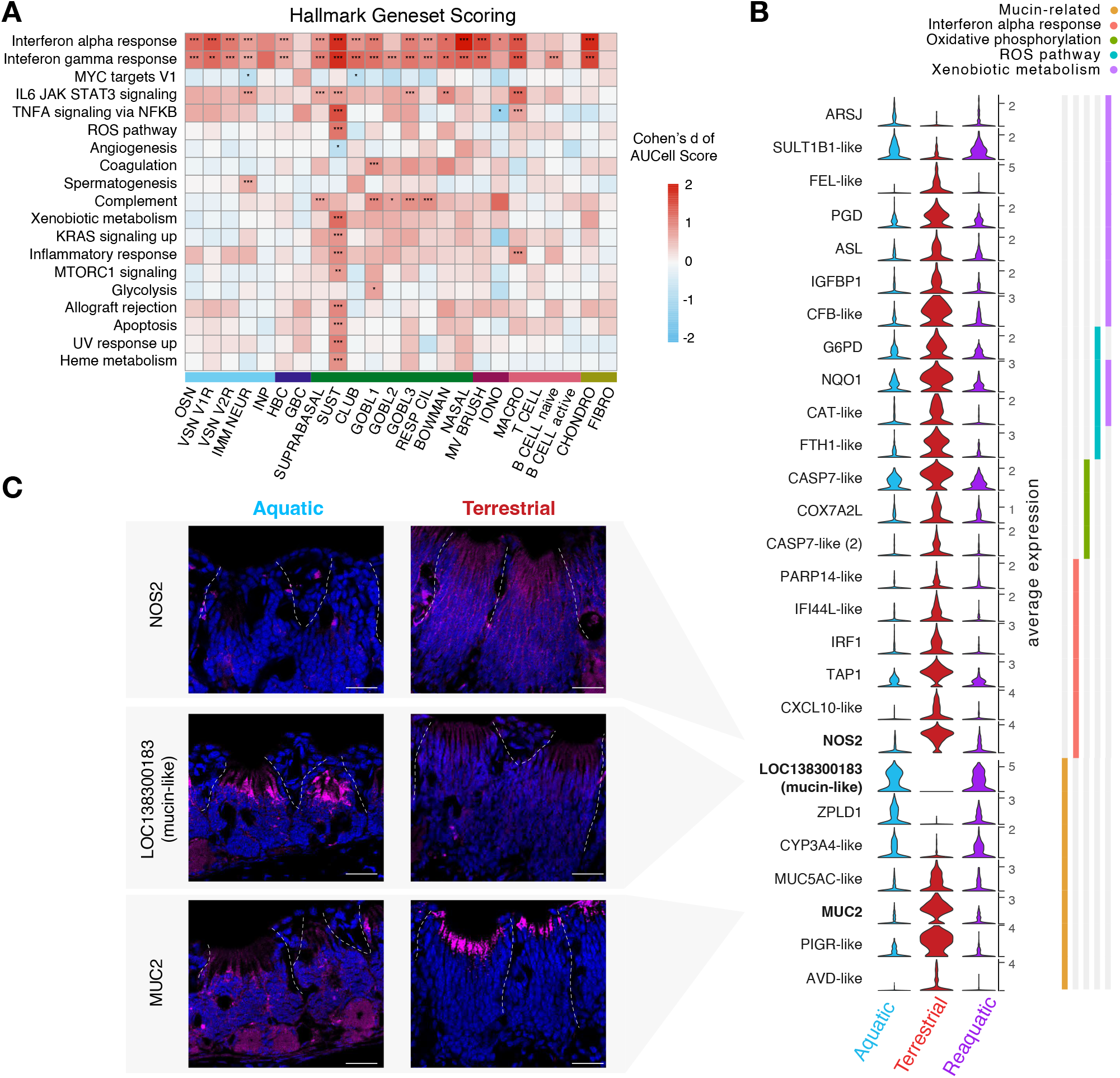
Transcriptomic plasticity of sustentacular cells across environments. A) Gene set enrichment analysis: for each cell type and for each gene set from the Hallmark50 set, the matrix shows the effect size (Cohen’s d) of gene expression enrichment (AUCell scores) between terrestrial and aquatic cell types. Only Hallmark gene sets with differential enrichment in at least one cell type are shown. Negative scores indicate the gene set is upregulated in cells from aquatic animals, while positive scores indicate the gene set is upregulated in cells from terrestrial animals. Significance determined by MAST algorithm, only effect sizes greater than 0.7 shown. B) Violin plots showing genes differentially expressed in sustentacular cells across the three conditions, grouped by functional category. C) Close ups on the olfactory epithelium showing differential expression of NOS2 (magenta), a mucin-like gene (LOC138300183), and the MUC2 gene, respectively, in sustentacular cells of terrestrial and aquatic animals. Scale bar 50um.

Transcriptomic plasticity was also observed in the three types of goblet cells (Supplemental Fig. 5, Supplemental Fig. 6). In both sustentacular and goblet cells, genes encoding mucins and other mucus-associated proteins are differentially expressed between environmental conditions (Supplemental Fig. 6). In sustentacular cells, for example, the gel-forming mucin MUC2 was upregulated in terrestrial conditions while a predicted mucin-like glycoprotein (LOC138300183) was upregulated in aquatic and re-aquatic conditions (Fig 5B,C). These observations indicate that the transition from an aquatic to a terrestrial lifestyle reformats the composition of the nasal mucus through transcriptomic changes in non-neuronal cells, possibly to cope with the higher biotic and abiotic stressors in the terrestrial environments (Wei et al., 2026). Changes in mucus composition may also influence odor detection; the diffusion of chemicals in the nasal cavity and their binding to sensory neurons depend heavily on the local extracellular milieu (Bryche et al., 2021, Shirai et al, 2023, Acquah et al., 2026). Taken together, these results demonstrate that gene expression in the olfactory system of the newt is plastic and reversible, just like morphology and locomotor behavior.

### Environmental change modulates neurogenesis

The odorant space available to animals is drastically different in water and on land. Fish and tetrapod genomes have different repertoires of chemoreceptors, and gamma-family olfactory receptors (ORs), which preferentially bind volatile chemicals, are greatly expanded in tetrapods (Niimura 2009, Policarpo et al., 2024). In frogs, metamorphosis coincides with the development of a new section of the olfactory epithelium called “air-nose” for its specialized role in detecting airborne odorants (Weiss et al., 2021, Reiss and Eisthen, 2008).

To test whether the transition to land upregulates the expression of receptors detecting volatile odors, such as gamma-family ORs, we compared the expression of chemoreceptors in aquatic and terrestrial *Pleurodeles*. We first annotated all chemosensory receptor genes in the *Pleurodeles* genome using an established pipeline (Policarpo et al. 2024) that assigns family identity to all ORs (1125 genes), trace amine receptors (TAARs, 10 genes), and vomeronasal receptors (V1Rs and V2Rs, 29 and 221 genes respectively, Supplemental Fig. 7A). Given the high number of chemoreceptor genes and their low detection rate in the scRNA-seq dataset, we turned to bulk RNA sequencing to assess their expression (n=4 animals for each condition). The large majority of chemoreceptor genes were detected in our bulk RNA-seq dataset (1282/1385), but we did not find any systematic difference in the expression of chemoreceptors at the family level (Fig. 6A), indicating that the transition from an aquatic to a terrestrial habitat does not trigger any global shift in gene expression from an “aquatic” to a “terrestrial” chemoreceptor gene repertoire.

**Figure 6:**
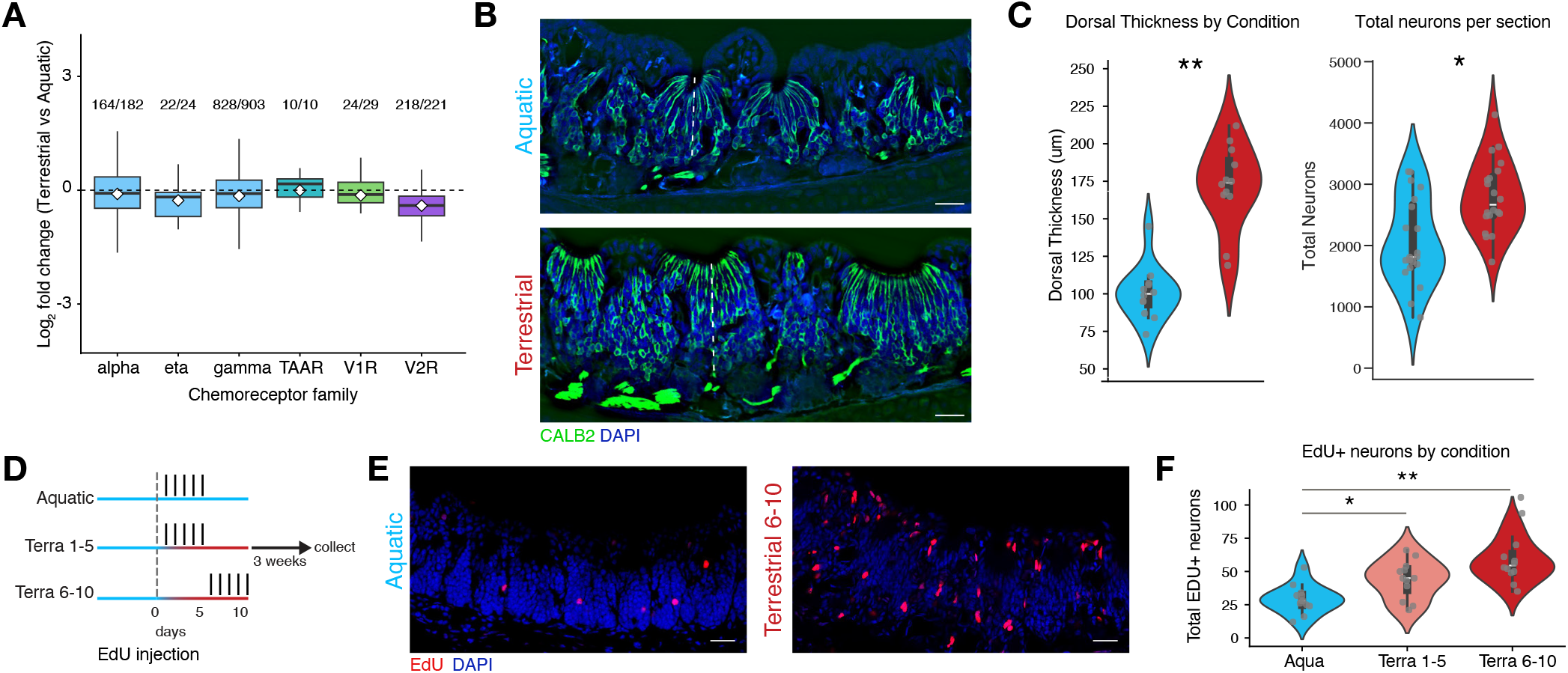
Increase of neurogenesis with the transition from an aquatic to a terrestrial habitat. A) Bulk RNAseq of nasal epithelia does not indicate differential expression of chemoreceptor gene families across conditions. Boxplots showing Log2 Fold change of chemoreceptor genes grouped by family, depicting mean (white diamond), median (thick black line), and interquartile range (box). Positive fold change indicates increase in the terrestrial condition. Numbers above each family indicate the number of detected chemoreceptors detected in bulk RNAseq data over the total number of family members identified in the genome. We found no significant enrichment of chemoreceptor families using competitive gene set testing (Limma package) while adjusting for inter-gene correlation. B) Sections of terrestrial and aquatic nasal epithelia, showing multiple stripes of olfactory epithelia stained with Calretitin (CALB2, green), a marker of sensory neurons. Dashed white line indicates thickness. Scale bar 200um. C) (left) Terrestrial animals have significantly thicker epithelia as measured by the distance from the nasal cavity to the basal lamina of the dorsal side of the epithelium of N=3 animals, n=4 slices/animal (p = 2.43e-08, general linear mixed effects model, dots represent individual slice measurements). (right) Terrestrial animals have significantly more neurons per section than aquatic animals (p=1.20e-4, negative binomial mixed effects model, see Methods for details on neuron counting). D) Schematic of EdU pulse chase experiment at the transition from aquatic to terrestrial life. E) Representative sections of the nasal epithelia in a terrestrial newt (EdU between 6 and 10 days) and in an aquatic control from the EdU pulse-chase experiment, EdU+ cells in red. F) Quantification of the number of EdU+ neurons in the EdU pulse-chase experiment at the time of terrestrialization. Significant increase of EdU+ neurons when labelling occurred in the first (p=2.07e-02), or second (p=1.345e-05) five days on land, compared to an aquatic control. Tukey adjusted p-values from the negative binomial mixed-effects model.

We next asked whether there were any other transcriptomic changes in OSNs and VSNs that may account for differences in olfaction between environments. After surveying the expression of ion channels, chemotransduction, calcium handling, synaptic transmission, and other genes related to neuronal function, we did not detect any significant difference between aquatic and terrestrial newts (Supplemental Fig. 8A). The few genes differentially expressed in aquatic and terrestrial sensory neurons and identified from the analysis of scRNAseq data (Fig. 4) belonged to the same functional categories differentially regulated in the rest of the nasal epithelium (Fig.5, Supplemental Fig. 8B). This analysis indicates that genes related to neuronal function generally maintain stable expression levels in sensory neurons across environments (Supplemental Fig. 8A).

Finally, we reasoned that an increase in the overall number of sensory neurons in the nasal epithelia may contribute to the higher responsiveness of terrestrial animals to olfactory stimuli on land. To test this hypothesis, we compared histological sections collected from the olfactory epithelia of terrestrial and aquatic age-matched siblings. Cell types of interest were identified with the markers CTIP2, which is expressed at high levels in immature neurons, and moderate levels in mature neurons (Supplemental Fig. 9B), and SOX2, expressed in sustentacular cells, close to the epithelial surface, and in stem cells at the base of the epithelium (Fig 4C). The olfactory epithelium of terrestrial animals was thicker on both the dorsal and ventral sides of the nasal cavity (Fig. 6B,C, Supplemental Fig. 9A), suggesting an increase in the overall number of sensory neurons. To confirm this hypothesis, we segmented individual cell bodies using Cellpose (Stringer et al., 2020) and quantified the number of neurons and sustentacular cells annotated from their anatomical position and the expression of CTIP2 or SOX2. This analysis revealed that the nasal epithelia of terrestrial animals had on average more sensory neurons compared to aquatic animals (Fig. 6C).

The presence of more OSNs in terrestrial animals suggests that the transition to land may trigger an increase in neurogenesis. To test this hypothesis, we performed pulse-chase experiments with the thymidine analog 5-Ethynyl-2′-deoxyuridine (EdU), which was administered to newts in either their first 5 days of life on land, or between days 6 and 10; aquatic age-matched animals served as controls (Fig. 6D). Analysis of EdU+ cells three weeks later revealed more EdU+ CTIP2+ cells in terrestrial animals compared to aquatic controls, indicating that consistent exposure to terrestrial conditions leads to an increase in the number of newly-generated sensory neurons (Fig 6E,F). We propose that the addition of new sensory neurons enables appropriate olfactory behavior on land, acting as a form of gain control for the system, which may help newts detect lower concentrations of odors in air compared to water.

## Discussion

Our study uncovers environmentally-induced plasticity across levels of biological organization in a vertebrate, the Iberian ribbed newt *Pleurodeles waltl*. We show that transitions between aquatic and terrestrial lifestyles elicit specific and reversible changes in gene expression, body morphology and animal behavior. These findings are consistent with observations in *Pleurodeles* and other newt species, where environment-dependent variation in tail morphology, physiology, and feeding behavior has been documented (Zamora-Camacho, 2023, Degani and Meerson, 2024, Brown 1984, Walters and Greenwald, 1977, Heiss and Vylder, 2016). Collectively, the available evidence suggests that environmentally-driven plasticity may be a general feature of newts, a subfamily of salamanders that originated about 60 million years ago (Stewart and Wiens, 2025). Our study thus establishes *Pleurodeles waltl* as a tractable laboratory model to study the mechanistic underpinnings of environmental plasticity in the nervous system.

Our cellular and molecular analysis of the nasal epithelium shows how environmental plasticity can be implemented at the cell type level. Secretory and support cells, in particular sustentacular cells, react to environmental changes with transcriptomic plasticity. Sustentacular cells activate an epithelial stress response, consistent with their role as epithelial barriers. Sensory neurons, conversely, maintain a stable expression of chemoreceptors but increase in number on land, through increased neurogenesis. We propose that these cell-type specific changes act together to enhance odor detection on land, where chemicals diffuse more rapidly: with more sensory neurons and a different mucus composition, the nose would be able to capture and sense odors more efficiently.

More broadly, this study uncovers three fundamental properties that set apart plasticity in newts from forms of plasticity described so far in other vertebrates.

First, environmental plasticity is a global phenomenon, spanning body morphology, sensory and motor behaviors, and multiple cell types in the nervous system. Consistent with this, metabolism and physiology are also modulated by transitions between water and land (Brown 1984, Walters and Greenwald 1977, Kristin and Gvozdik, 2014). These global changes indicate that the differences between aquatic and terrestrial environments are so profound as to require the adaptation of multiple organ systems and at different levels of biological organization. Previous studies suggest that hormones may act as systemic signals for plasticity in newts (Vellano et al., 1970, Singhas and Dent 1975). Whether environmental changes are detected by a central sensor and how global signaling is initiated remain open questions.

Second, we find that plasticity is elicited by a remarkably simple manipulation: transferring animals from an aquatic to a terrestrial habitat. This sharply distinguishes newt phenotypic plasticity from the forms of plasticity described in other vertebrates, such as songbirds, where plastic changes, including in the brain, are tied to seasonal cycles (Cornez et al, 2020, Clayton 1998, Alliende et al., 2017, Mohr et al., 2020). In the natural habitat where Iberian ribbed newts live, the availability of vernal pools is broadly seasonal, but fluctuations in rainfall can generate unpredictable dry periods. Consistently, our findings show that plasticity can be triggered on demand, regardless of the season, indicating the existence of a yet-unidentified mechanism that couples sensing of environmental changes and induction of plasticity.

Third, we show that environmentally-driven changes are reversible, i.e. they are bidirectional. Tail morphology, locomotor behavior, and gene expression in the nasal epithelium all revert to the aquatic state once land-habituated newts are returned to water. This clearly distinguishes newt phenotypic plasticity from developmental plasticity, a form of phenotypic plasticity that enables some vertebrate species to adapt to extreme environments. Environmental stressors during development can induce changes in metabolism, physiology, and even anatomical traits, but these adaptations are often irreversible (Denver et al., 1998). For example, bichir fishes raised in water or on land develop differences in their skeleton (Standen et al., 2014). In newts, however, plasticity takes place in post-metamorphic life stages and is reversible, indicating that these vertebrates can flexibly switch phenotypes throughout their lifetime. These observations suggest that environmental plasticity in newts builds on the unusual flexibility of development and life cycles in salamanders (Joven et al., 2019, Yun 2021).

The adaptations needed to thrive in water and on land are so profound that most species are either aquatic or terrestrial. Permanent transitions to terrestrial life, which took place multiple times in animal history, were associated with convergent innovations in animal genomes (Vermeij 2017, Wei et al., 2026). Interestingly, the gene families that convergently expanded with the permanent shift to terrestrial life - genes involved in detoxification, oxidative stress, and xenobiotic metabolism (Wei et al., 2026) - are also those upregulated in newts adapted to land. This suggests that environmental plasticity in newts may have evolved by adding a new layer of regulation to the same gene families that expanded with vertebrate terrestrialization. We propose that studying plasticity in vertebrates that evolved convergently the ability to live at the water-to-land interface can reveal how environmental constraints shape animal physiology and behavior. With its expanding neuroscience and genetic toolkit (Jaeger et al., 2025, Elewa et al., 2017, Hayashi et al., 2025), the Iberian ribbed newt *Pleurodeles* waltl now offers the opportunity to dissect how aquatic and terrestrial lifestyles constrain neural systems during the evolution of adaptive behaviors.

## Supporting information

All supplemental Figures and Tables

## Acknowledgements

We would like to thank J. Barber, S. Cook, S. Ngai and the Columbia University Institute for Comparative Medicine for animal care, all members of the Tosches lab, S. Lomvardas and L. Weiss for their critical feedback on the project, N.J. Chua for assistance with computational resources and data storage and management, V. Saltz, J. de la Cruz, V. Zhao, L.M.R Grajales, and L. Masoudi for pilot experiments and analyses, and A. J. Gillis for assistance with paraffin sections. Bulk RNA sequencing was conducted by the JP Sulzberger Columbia Genome Center, funded through the NIH/ NCI Cancer Center Support Grant P30CA013696. This work was supported by grants to MAT from the National Institute of Health (R35GM146973), McKnight Foundation, Chan Zuckerberg Initiative, Pershing Square Foundation, and Rita Allen Foundation. We also acknowledge support from the National Science Foundation (Graduate Research Fellowship to ECBJ), Revson Foundation (GG) and the University of Basel (to MP and WS).

## Author contributions

AMMM and MAT designed the project. AMMM and ECBJ performed behavioral experiments. AMMM and NMS collected animal morphometrics data. JW, AMMM, TGZ, GJG, and GG performed histology. JW and AOG generated sequencing data. MP annotated *Pleurodeles* chemoreceptors with input from WS. AMMM, LTG, and MAT performed bioinformatic analyses. AMMM, JW, and MAT analyzed the data. AMMM and MAT wrote the first draft of the manuscript with input from JW, all authors provided feedback on the final manuscript. MAT supervised the study and acquired funding.

## Materials and Methods

### Animals

Adult *Pleurodeles waltl* were obtained from a breeding colony established at Columbia University. Aquatic animals were maintained in an aquatics facility at 20◦C in tanks of 16L of water, where 4.5L are replaced daily through automated water changes. One to six animals were housed in each tank. Water is first purified through reverse osmosis, and then water chemistry is adjusted with Instant ocean to a conductivity of ∼900mA, and with sodium bicarbonate to a pH of ∼7.4. Terrestrial animals were maintained in an enclosed cage on a damp substrate (eco carpet) and misted with tank water for 5 seconds every 1 hour and 20 minutes using a MistKing system, ensuring that humidity levels remained around 80%. All animals were maintained on a 12L:12D cycle. All animals were fed a blend of bloodworms, agar and fish pellets 3x per week. All experiments were conducted in accordance with the NIH guidelines and the approval of the Columbia University Institutional Animal Care and Use Committee (IACUC protocol AC-AABL1150). Experiments were performed with adults (i.e. post-metamorphic animals) aged 1-4 years and all “steady state” terrestrial animals were continuously in terrestrial husbandry conditions for at least 6 weeks before use in experiments. All reaquatic animals had been returned to the water for at least 4 weeks after being on land before being used for experiments.

### Morphological analysis

For measurements of animals from the colony, newts were briefly anesthetized using MS-222 (0.2%), measured with digital calipers, and photographed before being returned to their home tanks. For time course experiments we measured unanesthetized animals to avoid submerging terrestrial animals repeatedly in liquid. To account for varying body sizes of animals in the colony, we normalized the width, height and length measurements by the snout-vent length. For significance testing we fit a linear model (statsmodels ols function in python) with environment, and mean-centered age as predictors for each trait. We first tested for an environment by age interaction, when the interaction was not significant we reported the additive model. P-values for the environmental effect were corrected for multiple comparisons using the Benjamini-Hochberg FDR procedure.

### Behavioral experiments

#### Wind tunnel experiments

Animals were tracked in a long, narrow acrylic arena (122 cm x 15 cm) lined with dampened paper sheets and white acrylic walls (21cm height, Canal Plastics, custom design) using a monochrome USB3.0 camera (Basler Ace2 a2A2590-60umBAS). A thin sheet of transparent acrylic was placed on top of the arena at the start of each trial to prevent odor from escaping. A custom olfactometer delivered either clean air or odor to the arena with rapid switching three way solenoid valves (LHDA1233115HA, Lee Company). The camera and stimulus timing were controlled and synchronized using custom Bonsai software and an Arduino Uno running Firmata. Air from the building source was controlled using a regulator (10psi - McMaster-Carr) and then passed through a charcoal filter. Airflow rate for each stream (wind or odor) was set by a flowmeter set to 1L/min. Each line could be diverted away from the arena to prevent pressure build up. Newts experienced 5 minutes of baseline (no stimulus), 5 minutes of clean air passed over a 20 mL scintillation vial filled with tank water, 5 minutes of odorized air where clean air passed over a 20mL scintillation vial filled with 15mL of crushed bloodworms (midge larvae) in tank water, followed by 5 more minutes of clean air over tank water, and one minute with no stimulation at the end.

Experimental protocol: One to two tanks of newts were run each experimental day, with 3-6 newts inhabiting each tank. Newts were food deprived for 6-14 days prior to the experiment. All newts were between 1-4 years old and we used males and females indiscriminately. At the beginning of each day newts were taken from their home tank and placed in a smaller transport container. Newts from terrestrial tanks were placed in containers on damp paper towels by carefully lifting them by their tails, aquatic newts were transferred carefully to the container using a net. Each trial lasted 21 minutes and newts ran 1-3 trials a day. Newts were individually identified by characteristic features (marks on the body, head spot pattern, size, and weight) or an RFID tag (Biomark) that was implanted into the tail or lower abdomen at least 1 month prior to the experiment. Newts were carefully positioned in the centre of the arena at a random orientation by picking them up by their tail and supporting them with the other hand to release them into the arena. Newts were returned to the transport container after each trial. At the end of the day newts were returned to their home tank. Some newts were run multiple times over the course of data collection.

#### Analysis of wind tunnel behavioral data

A deep learning network (supervised LightningPose model) was trained for keypoint annotation on 21 body parts (nose, left-side of head, right-side of head, neck, 3 points on the spine, the base of the tail, 3 points on the tail and the tail tip, and the left and right elbows, knees, feet and hands) from 780 frames from a variety of terrestrial and aquatic individuals using Lightning Pose (Biderman et al., 2024). Twenty random frames were selected per trial for annotation and context frames (two frames before and two frames after the selected frames) were also extracted. We trained the model using 500 training steps with ‘dlc-top-down’ as the base model using different random start seeds. We trained the model using the supervised-context settings in Lightning Pose. This data was imported as the basis for the Keypoint-MoSeq (Weinreb et al., 2024) model. We discarded the tail tip point at this stage as we found it had poor tracking. We trained Keypoint-MoSeq models to annotate behavioral syllables using a subset of well tracked data from the Lightning Pose model’s output. The training data was a blend of trials from terrestrial and aquatic animals. The initial AR (auto-regressive) model was trained using 8 principle components (to explain 90% of pose variation) for 50 iterations with a kappa value of 1e-8, and then the non-AR model was trained for an additional 450 iterations with a kappa value of 1e-7. This model produced syllables with an average duration of ∼400ms, similar to what has been validated in rodent studies (Weinreb et al., 2024)). We applied this model to the remainder of our data. Further analysis was completed using the syllables, centroid positional information and velocities generated by the Keypoint-MoSeq pipeline. The terrestrial-only model was constructed similarly but trained on selected trials where terrestrial animals went upwind towards the odor source. We adjusted the kappa parameter in Keypoint-MoSeq to obtain similar syllable lengths to our first model. We subsequently analysed the data with custom python scripts and the visualization functions available in the Keypoint-MoSeq package. Frequency was defined as in Weinreb et al., 2024 where it is the number of times a syllable initiates divided by the total number of syllable initiations.

To compute the trunk straightness index we calculated the distance from the neck keypoint to the base of the tail keypoint. We divided this by the sum of the lengths of the trunk segments (neck to spine1, spine1 to spine2, spine2 to spine3, spine3 to tail base). This index achieves a maximum value of 1 (perfectly straight). We took the average trunk straightness index for all frames where the animal was walking (defined by the walking syllables 0,1,2,3,4,5,6,7,12 in Fig. 2). To exclude brief walks, but allow for brief changes to other syllables/noise we found all instances where animals walked for >3s with <10 consecutive non-walking syllables. We calculated the average trunk straightness index, and then averaged this per trial.

When computing average head bend angle and angular velocity of the head, we filtered the Lightning Pose keypoint data for when the animal was facing the source (facing within 60 degrees of the source direction), reasoning that if they were walking away from the flow they were not actively engaging with it. We computed the angle between the vectors of the second keypoint on the spine to the neck, and the neck to the nose. We took the absolute value for joint angles so that bends to the left side and right side of the body were considered equal (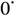 when nose and spine are aligned). We applied a median filter over 5 frames to smooth keypoint tracking jitter in these analyses. We then computed the median head bend angle and median angular velocity per trial..

#### Breathing rate behavior acquisition and analysis

To record the breathing rate of terrestrial and aquatic animals we placed them in a small acrylic box measuring (15cm x 15cm x 15cm) that had a small odor inlet, and was illuminated with IR LEDs from above and below. All experiments were performed in the dark and animals were habituated to a dark room for 30+ minutes before the experiments, and allowed to habituate to the box for an additional 10 minutes before the start of the trial. Video was acquired through 4 cameras (Basler ace2 a2A2590-60um-BAS (x3) and one allied vision alvium 1800U) that were controlled through custom bonsai software (Lopes et al., 2021), which also controlled stimulus output using an arduino Uno running Firmata. The cameras were synchronized by a hardware trigger, running at 31.25 frames per second, using a custom arduino script running on an arduino Mega. Wall air was passed through a charcoal filter, and split between a clean air flowmeter and an odor flowmeter, each set at 0.5L/s. Valves (LH-DA1233115HA, Lee Company) controlled the flow to two 20mL scintillation vials filled with tank water from the *Pleurodeles* colony (clean air) or 15mL of tank water with 2 cubes of frozen bloodworms dissolved in it. Flow was directed to a port on the chamber by a third valve. Synchronized video was imported in the BORIS software (Fraird and Gamba, 2016) and breaths were manually annotated as point events. This data was analyzed with custom python scripts.

#### Statistical analysis of behavioral data

The design of the gait and plume tracking experiments involved the same animals being run for multiple trials an unequal number of times. To account for this we used statistical models with repeated measures designs for this data. For syllable data between environmental conditions we averaged syllable frequencies within animals, and tested for differences in syllable frequency using a permutation-based Kruskal-Wallis test with Dunn’s post-hoc comparisons and Benjamini-Hochberg FDR correction (Weinreb et al., 2024). For the terrestrial model, wind and odor data were not independent groups as they were from the same animal at different points in time. To account for this we took the within-trial difference in syllable frequencies and fit a linear mixed effects model with stimulus as a fixed effect, and animal identity as a random intercept.

For the trunk straightness index we fit a linear mixed effects model with condition (terrestrial, aquatic, reaquatic) as a fixed effect and animal identity as a random intercept, and aqua as the reference level.

For mean arena position change during wind and odor, and mean occupancy of the upper portion of the arena, we calculated the difference between the average position in the second half of the wind and second half of the odor period, or the difference in occupancy. We then fit a linear mixed effects model with a fixed effect of condition (terrestrial or aquatic) and a random intercept of animal identity.

For head bend angle and angular velocity, we fit a linear mixed effects model as above, where there was a fixed effect of stimulus (wind or odor) and a random intercept of animal identity. In all cases we corrected for multiple comparisons using the Benjamini-Hochberg FDR procedure.

The kruskal-wallis test was implemented via the Keypoint MoSeq package, and all other linear mixed effects models were fit with statsmodels mixedlm function in python (3.9.13 for arena position and occupancy, 3.14.4 for head bend analysis and straightness index)

### EdU administration

EdU was injected intraperitoneally (50mg/kg of total body weight, made from a 5mg/mL stock solution in 0.9% saline) using an insulin syringe. For the neurogenesis experiments, animals were injected once daily at the same time of day for 5 consecutive days (starting 1 day, or 6 days after beginning to live on land).

### Histology

#### Tissue preparation

Animals were anaesthetized by submersion in 0.2% MS-222 and perfused with 1x molecular grade PBS followed by 4% PFA in 1x molecular grade PBS intracardially, before dissecting the snout from the head. Snouts were split in half, to isolate individual epithelium. This tissue was fixed overnight with 4% PFA in molecular grade PBS.

#### In situ hybridization

Following fixation, the tissue was dehydrated by increasing gradient of methanol in PBS. Tissue was stored at -20 C in 100% methanol until further processing. Snouts were bleached overnight in 5% H2O2 in methanol. Whole mount tissue was washed in Probe Wash Buffer (Molecular Instruments) 2 x 30 min, then prehybridized in Hybridization Buffer (Molecular Instruments) at 37°C for 1 hour. Probe pairs were designed using the insitu_probe_generator (Kuehn et al., 2021) and ordered from IDT. Tissue was stained by incubation in 2-6 pmol of each probe set (Probe sets available in supplementary file) for 3 nights at 37◦C, washed 3 x 45 minutes in Probe Wash Buffer (Molecular Instruments) at 37◦C, followed by 3 x 45 minutes washes in 5xSSCT at room temperature (RT). The tissue was then incubated in Amplification Buffer (Molecular Instruments) for 2 hours at 4◦C, then incubated at 4◦C for 3 nights in snap-cooled hairpin solution (30 pmol each, Molecular Instruments). Following amplification, the tissue was washed 2 x 1 hour in 5xSSCT, then 2 x 1 hour in 500 mM Tris-HCl. Snouts were decalcified in 10% EDTA in 0.1 M Tris pH 7.2 for 3 nights at 4◦C, embedded in 4% agarose in 500 mM Tris-HCl, and 100 um coronal sections collected using a Leica VT1200S vibratome. Sections were DAPI stained 2 hours at RT in Tris-HCl, then mounted in DAKO fluorescent mounting medium (Agilent Technologies). Images were acquired using a confocal microscope (Zeiss LSM800) and processed in FIJI (Schindelin et al., 2012).

#### Histochemical staining

Fixed snouts from aquatic and terrestrial salamanders were washed twice with PBS and decalcified in a solution of 10% EDTA in 0.1M Tris (pH 7.2) for 2 days at room temperature. Tissue preparation and staining was then performed following Rees et al (2024). Snouts were separated in left and right halves, cleared with Histosol (National Diagnostics) 3×20 min at room temperature, moved into a solution of 1:1 Histosol:paraffin for 2×30 min at 60°C, and then infiltrated with molten paraffin overnight at 60°C. After an additional 3×1 hour paraffin changes, samples were embedded in square Peel-A-Way molds (Sigma-Aldrich), left to set for 24 hours and then sectioned at 8 µm on a Leica RM2125 rotary microtome. Sections were mounted serially on SuperFrost Plus charged glass slides and processed for Masson’s Trichrome or Toluidine Blue staining. Histochemical staining with modified Masson’s Trichrome was performed as described in Witten and Hall (2003) using Mayer’s acid haematoxylin (Sigma MSH32), xylidene ponceau (0.5% xylidine ponceau in 1% acetic acid and 0.5% acid fuchsin in 1% acetic), 1% phospho-molybdic acid, and 2% light green in 2% citric acid (diluted 1:10 in water before use). For Toluidine Blue staining, dewaxed and rehydrated sections were incubated for 4 minutes in 0.4% toluidine blue in milliQ water (Himedia GRM1000) and counter-stained with 0.02% Fast Green in milliQ water for 3 minutes (Sigma-Aldrich F7252).

#### Immunohistochemistry

Tissue was washed to stop fixation 2×15 minutes in PBS, then decalcified for 3 nights in 10% EDTA in 0.1 M Tris-HCl pH 7.2. After agarose embedding, sections (100um) were obtained using a Leica VT1200S vibratome, then blocked in Blocking Buffer (2.5% BSA, 2.5% sheep serum, 50 mM glycine) in PBST (PBS with 0.2% Triton), then incubated with the primary antibody solution in 10 mM glycine, 0.1% H2O2 in PBST for 1-3 nights. One antibody (CTIP2, abcam: ab18465) required antigen retrieval. In these samples, before blocking, sections were permeabilized in PBS-Triton 0.2% for 30 minutes, and then incubated in 10 mM sodium citrate pH=6.0 for 30 minutes at 70◦C, and finally washed 3 x 5 minutes in PBS-Triton 0.2%. EdU was detected using the Click-IT reaction kit (Invitrogen), according to the manufacturer instructions. Slices were washed 3x 15 minutes in PBST, then incubated with DAPI and secondary antibodies conjugated to Alexa fluorophores for 2 hours at RT. Sections were mounted in DAKO fluorescent mounting medium (Agilent Technologies). Images were acquired using a confocal microscope (Zeiss LSM800) and processed in FIJI.

Animals used for olfactory epithelium thickness measurements and neuron counts were paired siblings (average age: 30 months (15-46 months), average time terrestrial: 16 months (4-30 months)) as were the animals used for the EdU neurogenesis experiments (12 months of age at time of terrestrialization, collected 3 weeks after environmental transition).

### Cell segmentation, counting, and classification

For the experiments in Fig. 6 we measured the thickness of the epithelia on both the dorsal and ventral sides from sections matched along the A-P axis from age matched siblings raised across different conditions using FIJI. We measured from the base of the SOX2+ nucleus (sustentacular cells) to the midpoint of the CTIP2+ layer. On a small subset of samples the entire slice could not be captured in one Z-plane. In these cases we took two sections to capture the entirety of the tissue and max projected the channels.

To count neurons across slices we used Cellpose to segment cell bodies from the DAPI channel. We manually drew ROIs to highlight the main olfactory epithelium, and exclude the respiratory ciliated cells close to the nasal surface, the VNO, the secretory epithelium, the glands and other tissue past the nasal cavity. Using custom python scripts we set thresholds on the fluorescence intensity to roughly classify cells as either mature neurons (moderate CTIP2, low SOX2), immature neurons (high CTIP2, low SOX2), sustentacular cells (high SOX2, low CTIP2) or None (low SOX2, low CTIP2). Following this we used custom python scripts and Napari (Chiu et al., 2022) to correct annotations by hand. This involved: removing SOX2+ cells away from the outer nasal cavity (HBC/GBC cells), removing misannotated cells due to autofluorescence or other artifacts (including stripes of respiratory epithelium misannotated as OSNs), and removing any cellpose segmentation masks that were not truly cells. In the end we found the CTIP2 staining levels to be variable, and merged mature and immature neurons for further analysis after counting.

To count EdU+ neurons in the neurogenesis experiments we performed the same staining, but the lower number of cells allowed us to count and classify cells by hand using FIJI.

### Statistical analysis of histological data

In both the thickness, total neuron counts and EdU+ neuron counts we sampled 4 sections per animal along the A-P axis, which were matched across animals. For thickness we used a linear mixed effects model, with condition and slice position as fixed effects and animal identity as a random intercept. This was implemented with statsmodels mixedlm function in python. Since the total number of neurons and number of EdU+ cells were count data, we used R’s GlmmTMB package to fit a negative binomial generalized linear mixed model with condition and section level were fixed effects and animal identity was a random intercept. For the EdU+ neuron counts we used the emmeans package to perform pairwise comparisons on the estimated marginal means derived from the mixed effects model, using Tukey’s HSD to control for family wise error rate across contrasts.

### Single-cell RNA sequencing experiments

#### Epithelia dissociation and single cell capture

Terrestrial, aquatic and reaquatic animals were collected for epithelial dissociation, with two biological replicates for each condition.

Two pairs of siblings were used for the Terra and Aqua animals (32 months and 18 months old). The 32 month old terrestrial animal lived on land for 16 months, while the 18 month old terrestrial animal lived on land for 9 months. Reaquatic animals were siblings, 38 months of age, lived on land for 25 months before being returned to water for 3 months. We matched sex across conditions, so each condition had one male and one female representative. Animals weighed 4.5-10g, averaging 8g. Sibling animals from each condition were anesthetized by submersion in 0.2% Ethyl 3-aminobenzoate methanesulfonate (MS-222) before being perfused transcardially with cold, carbogenated, calcium-free Amphibian Ringer solution (96 mM NaCl; 20 mM NaHCO3; 2 mM KCl; 10 mM HEPES; 11 mM glucose; 0.5 mM MgCl2) and then decapitated. Snouts were transected at the olfactory nerve level, the nasal cavity was dissected open, and the nasal epithelium, including parts of the underlying palate, was removed carefully and cut into cubes in carbogenated Ringer. All tubes were coated with 1% BSA in HA -Ca media (Hibernate A, Calcium free, Brain Bits). The tissue was transferred to a 5 mL tube in 2.5mL of dissociation buffer (TTX, Liberase, DNase, and Papain in TrypLE) and incubated at RT for 30 minutes, and shaken head-to-tail on a rotator. Following the enzymatic dissociation, the tissue was mechanically dissociated on ice by trituration with fire polished, silanized glass pipettes with decreasing tip diameter. The supernatant was transferred to a new tube, while the cartilage was rinsed and discarded. The suspension was passed through a dampened 100 um cell strainer into a 50 mL tube, and additional media was added up to 20mL. The suspension was then passed through a dampened 70 um cell strainer, and a 5 mL BSA cushion (4% bovine serum albumin (BSA) in calcium-free, phenol red-free Hibernate) was added to the bottom of the tube, providing a density gradient to filter debris. The cells were centrifuged at (300xg at 4◦C) for 5 minutes, the supernatant was removed, and the pellet was resuspended in 100-180 uL HA -Ca -Mg media (calcium-free, magnesium-free, BrainBits). The suspension was passed through a 40 um filter into a 1.5 mL tube, and resuspended by pipetting up and down gently. The cell concentration was determined by counting on a Fuchs-Rosenthal chamber with trypan blue, and diluted to 1000 cells/uL in HA - Ca -Mg. The cells were then loaded onto a 10x Chromium Chip G for GEM generation and cell barcoding, with one reaction per condition, each targeting a cell recovery of 8000 cells. Single-Cell RNA-seq libraries were prepared with the 10x Chromium Next GEM Single Cell 3’ Reagent kit v3.1. and sequenced with NovaSeq X Plus (PE150) averaging 130480 reads per cell, and averaging 3309 UMIs per cell after removing empty droplets but before quality filtering and 8361 after filtering.

### Annotation of *Pleurodeles* chemoreceptors and gene models update

The *Pleurodeles* genome had limited categorization and information on chemosensory receptor genes, and many genes named by their locus identifier, which limited biological interpretation. We used a multi-pronged approach to identify previously unannotated chemoreceptor genes, improve gene models for chemoreceptors and other genes expressed in the nasal epithelium, assign family information to the chemoreceptor genes, and assign HGNC symbols for locus-identified genes. First, we performed full-length mRNA sequencing (PacBio IsoSeq 3) of a library generated after pooling RNA extracted from the nasal epithelia of 3 terrestrial and 3 aquatic animals; these IsoSeq data were used to refine gene models, to improve mapping to genes expressed in the nasal epithelia. Beginning with the gene transfer format file (GTF) from the NCBI genome available at (https://ftp.ncbi.nlm.nih.gov/genomes/all/GCF/031/143/425/GCF_031143425.1_aPleWal1.hap1.20221129/), we retained the longest gene model between the IsoSeq and published genome. Next, we added additional gene models for chemosensory genes identified through homology as described in Policarpo et al., 2024 as well as the mitochondrial genes. In addition, each chemoreceptor was assigned to a subfamily, according to the classifications in Policarpo et al., 2024. To improve biological interpretability we also created a mapping table to estimate the most similar HGNC named gene, and replaced locus identified genes with these for presentation purposes. We did this by a combination of text-based matching on protein and gene names to the Human, mouse, and Xenopus laevis genomes. We downloaded the genome annotation for all *Pleurodeles* genes for the GCF_031143425.1_aPleWal1.hap1.20221129 genome and subset all symbols beginning with ‘LOC’ (the locus identifier genes) for protein coding genes. Next, all genes labelled ‘uncharacterized protein’ were removed from further analysis. We stripped the ‘-like’ suffix from these genes and string matched on the ‘Name’ column, followed by the ‘Protein Name’ column of the tables, in the order of human, xenopus, mouse. Once a gene had matched, it was excluded from further matching attempts. Remaining genes were matched using the eggNOG orthology mapper (Cantalapiedra et al, 2021). All matches not in the HGNC format were converted using the MyGeneInfo python package (Xin et al., 2016). Not all locus identified genes were matched to an HGNC format gene. All genes in the figures with the ‘-like’ suffix are locus identified genes that have been renamed for presentation purposes.

### Quality control and filtering of single-cell RNA sequencing data

Single-cell libraries were aligned using CellRanger v9.0.0. Gene matrices were loaded into the R package Seurat 5.0.1. Each condition was preprocessed independently to filter out cells with less than a minimum number of features (<700), or higher than 10% mitochondrial genes. We then merged the libraries and used Seurat’s SCTransform function to normalize the raw data, regressing by percentage of mitochondrial gene expression, G2M score and S score. We computed the top 3000 variable genes to use for downstream analysis. Next, the data were integrated using Seurat’s integration pipeline. Using SelectIntegrationFeatures function we selected the top 3000 genes that were variable across conditions and prepared the datasets using these genes. Anchors, defined as the pairs of mutual nearest neighbours were found using Seurat’s FindIntegrationAnchors with a ‘cca’ reduction (canonical correlation analysis) and the normalization method set to SCT. We integrated the data with the IntegrateData function in Seurat using the anchors we computed and 100 dimensions. For downstream analysis we performed hierarchical tree clustering and generated the UMAP embedding using CHOIR (Sant et al. 2025) on the integrated layer of the data. This resulted in a dataset with 15,813 cells (4582 terrestrial, 5244 aquatic and 5987 reaquatic) and 64 clusters. This integrated dataset was used to annotate cell types using marker genes from the mouse, human, and zebrafish literature.

We reasoned that the integration may have been removing a large portion of real biological variance between the libraries. Therefore we performed the same quality control metrics described above and then merged the data rather than integrating. We performed normalization using SCtransform and regressed by mitochondrial gene percentage, environmental condition, original library, G2M and S score, to remove batch-specific technical artifacts. This generated an object with the identical cells to our integrated object but with different normalization and embedding. We then transferred our broad class annotations from the integrated object to the merged object. We further curated this dataset by removing the cell types outside of the nasal epithelium (teeth, neuromasts, and hair cells). Next, we performed Louvain clustering in Seurat with a resolution of 1 and generated the UMAP embedding from the top 100 principle components. Two clusters with low feature counts and very low percentages of mitochondrial genes were then removed, before repeating the normalization and clustering procedures. This produced a final merged object with 14,739 single cell transcriptomes for further analysis. We noticed strong expression of FOXI1 and CFTR, classic ionocyte markers (Deprez et al., 2020), in a subset of support cells that was not clustering independently in the integrated data. We subset the merged object for only support cell classes (HBC, GBC, SUST, SUPRABASAL, RESP_CIL, CLUB, GOBL1, GOBL2, GOBL3) and reclustered the data with Seurat’s Louvian clustering (resolution=1). Here the ionocytes formed their own cluster, and we applied this cell type label back to the same cells in the merged object.

### Differential expression analysis

We used Seurat’s FindMarkers function to compute the differentially expressed genes (DEGs) across the dataset (merge Seurat object). Genes were considered differentially expressed if the Log fold change of their expression levels across conditions was > 1.5, with an adjusted p-value of 0.01, a minimum difference of cells expressing a gene in a cluster of 20%. We used the MAST algorithm in the FindMarkers function for the identification of DEGs with animal identity as a latent variable to control library-specific batch effects. As MAST poorly handles genes with all or nothing expression between conditions (such as LOC138300183, which we verified histologically), we included all genes that had true zero expression in one condition, and were significant using the Wilcoxon method as well. All DEGs were identified using the RNA layer of the object.

### Geneset Scoring, Overrepresentation analysis and functional categorizations

To functionally annotate the transcriptomic changes between environments at the cell-type level, we used the AUCell algorithm (Aibar et al., 2017) implemented by SeuratExtend’s GeneSetAnalysis function (Hua et al., 2025). We used the Hallmark50 genesets (Liberzon et al., 2015). Due to the large number of *Pleurodeles* specific duplications, we took the human Hallmark50 set, and identified all genes not present in our object. If these genes had an HGNC symbol match using our mapping table described above, we replaced the human gene with the Locus gene(s) (for instance if MUC5B was in a human gene set, it would be replaced with the multiple locus genes that were annotated as ‘MUC5B-like’). After performing AU-Cell scoring, we computed the average scores per condition, and calculated the effect size with Cohen’s d. We used the MAST algorithm’s hurdle model to test for significance, and adjusted p-values for multiple comparisons using the Benjamini-Hochberg FDR correction. We did not display significance values for |cohen’s d| <0.7.

To determine whether genes involved in neuronal function were differentially expressed in sensory neurons, we selected relevant gene sets from the KEGG, REACTOME and GO databases via the msigdbr package (Subramanian et al., 2005, Dolgalev 2026). We replaced the genes with *Pleurodeles* specific genes as described above. Specifically we used the following categories: olfactory transduction: KEGG_LEGACY (C2) OLFACTORY_TRANSDUCTION, Neuronal Function: REACTOME (C2) REACTOME_NEURONAL_SYSTEM, REACTOME_ION_CHANNEL_TRANSPORT, REACTOME_TRANSMISSION_ACROSS_CHEMICAL_SYNAPSES, “REACTOME_NEUROTRANSMITTER_RECEPTORS_AND_POSTSYNAPTIC_SIGNAL_TRANSMISSION, REACTOME_POTASSIUM_CHANNELS, REACTOME_VOLTAGE_GATED_POTASSIUM_CHANNELS, Action potential: from GO (c5) GOBP_ACTION_POTENTIAL, GOBP_NEURONAL_ACTION_POTENTIAL, GOBP_MEMBRANE_DEPOLARIZATION_DURING_ACTION_POTENTIAL, Calcium Handling: GOBP_CALCIUM_ION_TRANSPORT, GOBP_RELEASE_OF_SEQUESTERED_CALCIUM_ION_INTO_CYTOSOL, GOBP_CALCIUM_ION_REGULATED_EXOCYTOSIS. Synaptic transmission: GOBP_SYNAPTIC_SIGNALING. To ensure we did not miss any ion channels, we downloaded all ion channel genes from the HGNC gene names database (https://www.genenames.org/data/genegroup/#!/group/177) and replaced genes not found in our dataset with *Pleurodeles* specific locus genes as above. We took the union of all gene sets, as many genes share membership, and annotated with the following priority: Action potential, Ion channels, Olfactory transduction, Calcium handling, Neuronal systems, and synaptic transmission with earlier categories taking priority.

To identify mucins and related genes in *Pleurodeles* genome, we searched the NCBI genome annotation file for the following terms: “mucin”,’mucus’,’mucosal’,”glycoprotein”, “immunoglobulin”,”lectin”. Additionally we added the genes associated to the following GO terms: “GOCC_MUCUS_LAYER”, “GOBP_MUCUS_SECRETION”, “GOBP_POSITIVE_REGULATION_OF_MUCUS_PRODUCTION”, from the msigdbr package. Finally, we added a small number of custom genes we determined could be involved in the mucus composition by examining the DEG lists of support cell classes.

### Overrepresentation analysis

To perform overrepresentation analysis on the DEGs of sensory neurons, we selected the OSN DEGs as described previously, and used the enricher function in clusterprofiler (Wu et al., 2021). We set the universe to all other genes which were expressed in at least 20% of sensory neurons in at least one condition, and used the Benjamini-Hochberg False discovery rate procedure to correct for multiple comparisons. We used the C5 collection of human gene sets from msigdbr to compute enrichment against all gene ontology sets (Biological processes, molecular function and cellular compartment), and replaced genes with *Pleurodeles* specific genes as described previously. We used an adjusted p-value cutoff of 0.05, and a minimum gene set size of 10 and a maximum of 500.

### Single-cell Data representation

All plots were created using either Seurat’s default plotting functions, SeuratExtend’s plotting functions, the ggplot2 package or Clusterprofiler’s plotting tools.

### Bulk RNA sequencing experiments

Total RNA was extracted from the epithelia of four animals living in an aquatic environment, and four animals living in a terrestrial environment. Animals were 14-37 months of age (21 months average), and terrestrial animals were on land 6-15 months (12 months on average). Animals were perfused with Ringer’s solution, and the epithelium dissected out. The tissue was homogenized and cells lysed with Trizol by mechanical dissociation. Chloroform was added, and the solution was shaken and centrifuged 15 minutes at 12000 rpm at 4°C to separate the RNA into the aqueous phase, discarding DNA, proteins, and lipids. Additional organic extractions were performed with the addition of chloroform or phenol:chloroform:isoamyl alcohol, while taking only the aqueous phase to remove contaminants. Quality check of the RNA was performed on the Agilent Bioanalyzer instrument, using the Eukaryote Total RNA Nano kit to determine RIN and concentration. Only samples with RIN values over 8 were included for bulk sequencing. Library preparation and sequencing were carried out by the Columbia Core Sulzberger Genome Center on the AVITI platform to a targeted depth of 80 million reads per sample. Reads were aligned to a reference transcriptome made from our Isoseq-updated *Pleurodeles* genome using Kallisto (Bray et al., 2016). We filtered out all genes with less than 10 total counts across all samples. We performed differential expression analysis using the DESEQ2 package (Love, 2014), contrasting terrestrial and aquatic environments. To test for statistical significance at the family level we used edgeR’s calcNormFactors to normalize raw counts and transformed using voom (Law et al., 2014) Statistical analysis was performed using the competitive gene-set testing for each chemoreceptor family using the camera function in the limma package (Wu and Smyth 2012, Ritchie et al., 2015), with inter-gene correlation empirically estimated for each family using the interGeneCorrelation function, rather than the default because of the high co-regulation among chemoreceptor genes. P-values were corrected for multiple comparisons with the Benjamini-Hochberg FDR procedure.

## Notes

### Competing Interest Statement

The authors have declared no competing interest.

