## Supplementary material for "Cell-type Plasticity Supports Behavioral Adaptations at the Water-to-Land Interface": All supplemental Figures and Tables

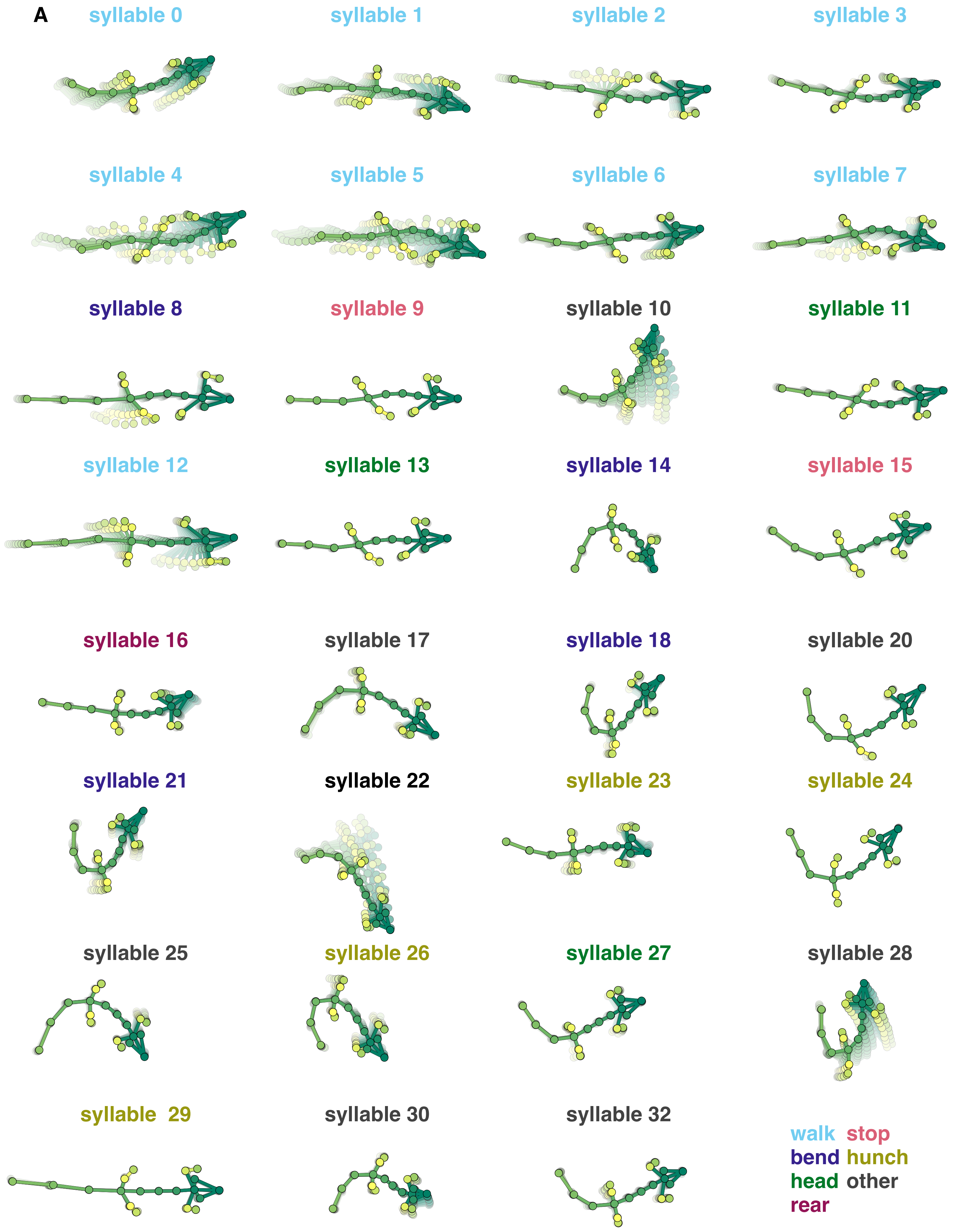
A
syllable 0
syllable 1
syllable 2
syllable 3
syllable 4
syllable 5
syllable 6
syllable 7
syllable 8
syllable 9
syllable 10
syllable 11
syllable 12
syllable 13
syllable 14
syllable 15
syllable 16
syllable 17
syllable 18
syllable 20
syllable 21
syllable 22
syllable 23
syllable 24
syllable 25
syllable 26
syllable 27
syllable 28
syllable 29
syllable 30
syllable 32
walk stop
bend hunch
head other
rear

**Supplementary Figure 1: Behavioral syllable repertoire in aquatic and terrestrial newts moving on a terrestrial substrate, related to Figure 2**

Average trajectories (keypoint change over time) of behavioral syllables identified by a Keypoint-MoSeq model trained on terrestrial and aquatic locomotion data, color indicates manually annotated behavioral category.

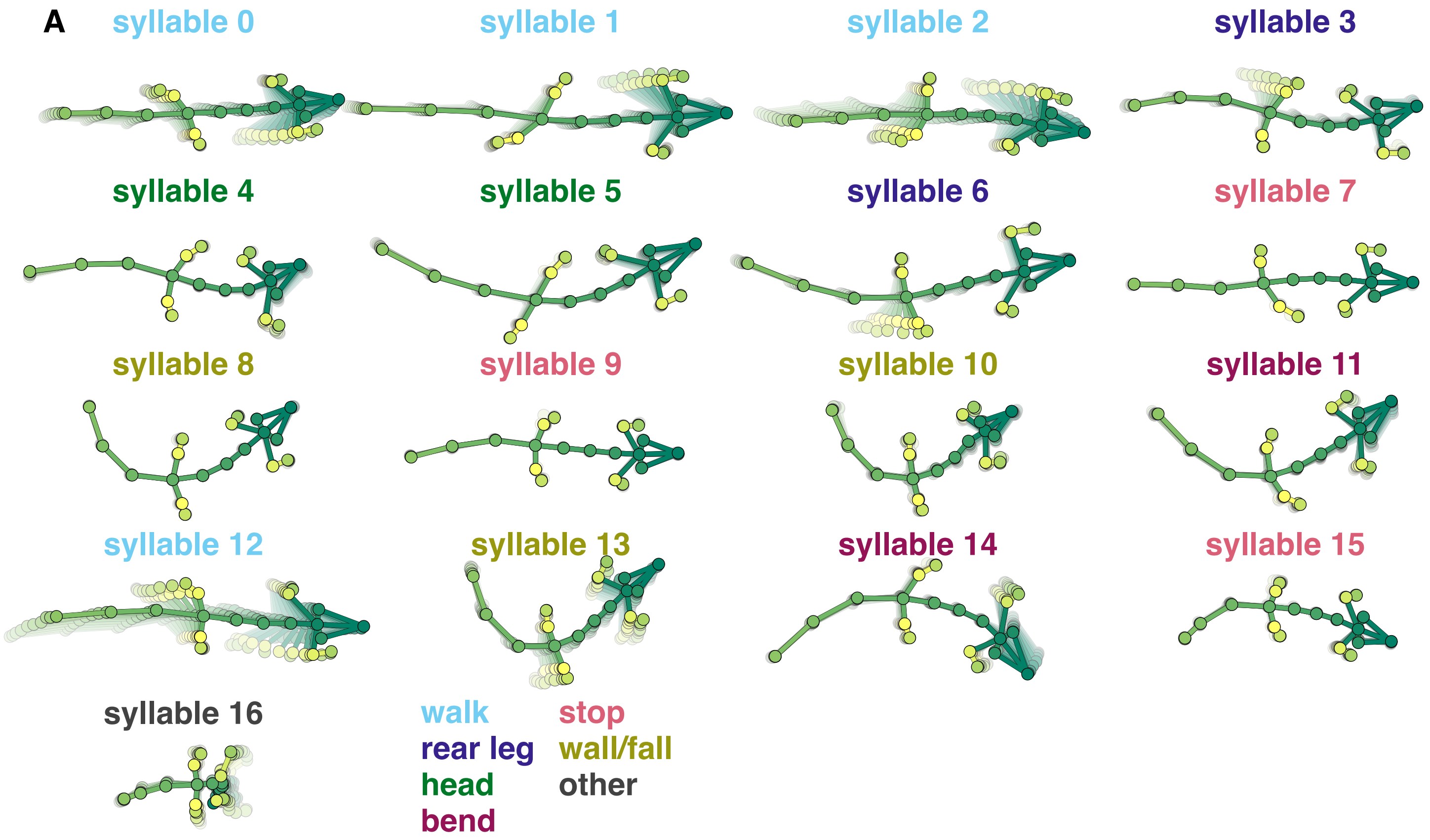
A
syllable 0
syllable 1
syllable 2
syllable 3
syllable 4
syllable 5
syllable 6
syllable 7
syllable 8
syllable 9
syllable 10
syllable 11
syllable 12
syllable 13
syllable 14
syllable 15
syllable 16
walk
stop
rear leg
wall/fall
head
other
bend

**Supplementary Figure 2: Behavioral syllable repertoire in terrestrial newts during olfactory navigation, related to Figure 3**

Average trajectories (keypoint change over time) of behavioral syllables identified by a Keypoint-MoSeq model trained on terrestrial data only, where animals moved towards the odor source, color indicates manually annotated behavioral categories.

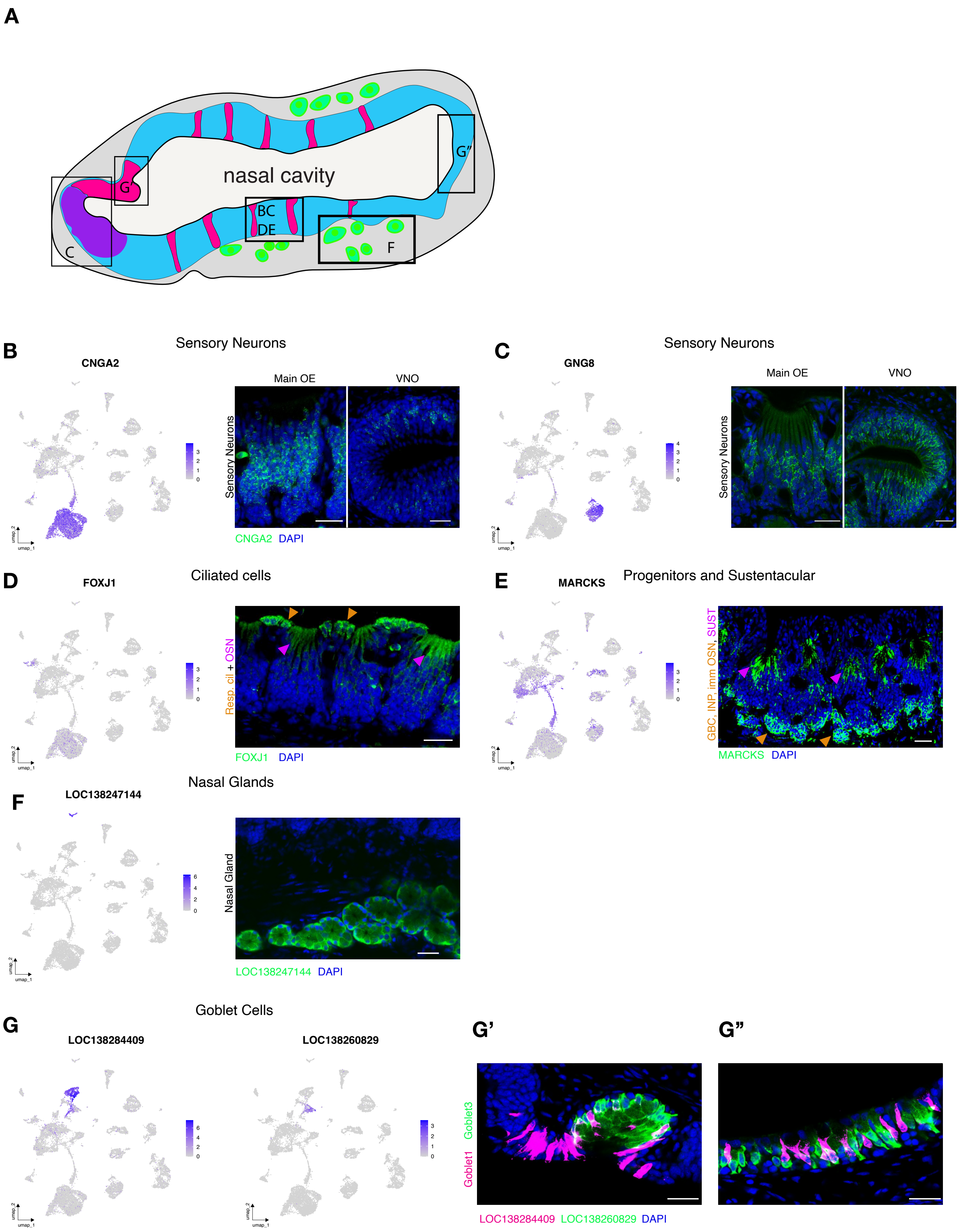

A
nasal cavity
G'
G''
BC
DE
C
F
B
CNGA2
Sensory Neurons
Main OE
VNO
Sensory Neurons
CNGA2 DAPI
C
GNG8
Sensory Neurons
Main OE
VNO
Sensory Neurons
D
FOXJ1
Ciliated cells
Resp. cil + OSN
FOXJ1 DAPI
E
MARCKS
Progenitors and Sustentacular
GBC, INP, imm OSN, SUST
MARCKS DAPI
F
LOC138247144
Nasal Glands
Nasal Gland
LOC138247144 DAPI
G
Goblet Cells
LOC138284409
LOC138260829
G'
G''
Goblet3
Goblet1
LOC138284409 LOC138260829 DAPI

**Supplementary Figure 3: Cell type specific marker genes and their expression in the newt nasal epithelium, related to Figure 4**

A) Schematic of a coronal section of the nasal epithelium. Inset boxes with letters show where additional figure panels are located.
B) (left) UMAP showing CNGA2 expression, a marker of OSNs; (right) HCR of CNGA2 (green) labels sensory neurons in the main olfactory epithelium and in the VNO.
C) (left) UMAP showing GNG8 expression in V2R-expressing VSNs; (right) HCR for GNG8 (green) V2R VSNs in both the main olfactory epithelia and the VNO.
D) (left) UMAP showing MARCKS expression in GBCs, INPs, Immature sensory neurons and in sustentacular cells. (right) HCR for MARCKS (green) highlighting sustentacular cells (near nasal cavity, orange arrows) and GBC/immature sensory neurons/INPs near the basal lamina (magenta arrows).
E) (left) UMAP showing FOXJ1 expression in respiratory ciliated cells, and sensory neurons. (right) HCR for FOXJ1 (green) labels respiratory ciliated cells (bordering the nasal cavity, orange arrows) and sensory neurons(within OE stripes, magenta arrows).
F) (left) UMAP showing the gene LOC138217144 (5S ribosomal RNA), expressed exclusively in nasal glands. (right) HCR for LOC138217144 (green) in a population of glands.
G) (left) UMAPs showing the expression of LOC138284409 (MUC5B-like) and LOC138260829 (Ranaspumin-like), which mark the GOBL1 and GOBL3 clusters respectively. (right) Double HCR for LOC138284409 (magenta) and LOC138260829 (green) showing the goblet cells in the secretory epithelium. All scale bars 50um.

**A**

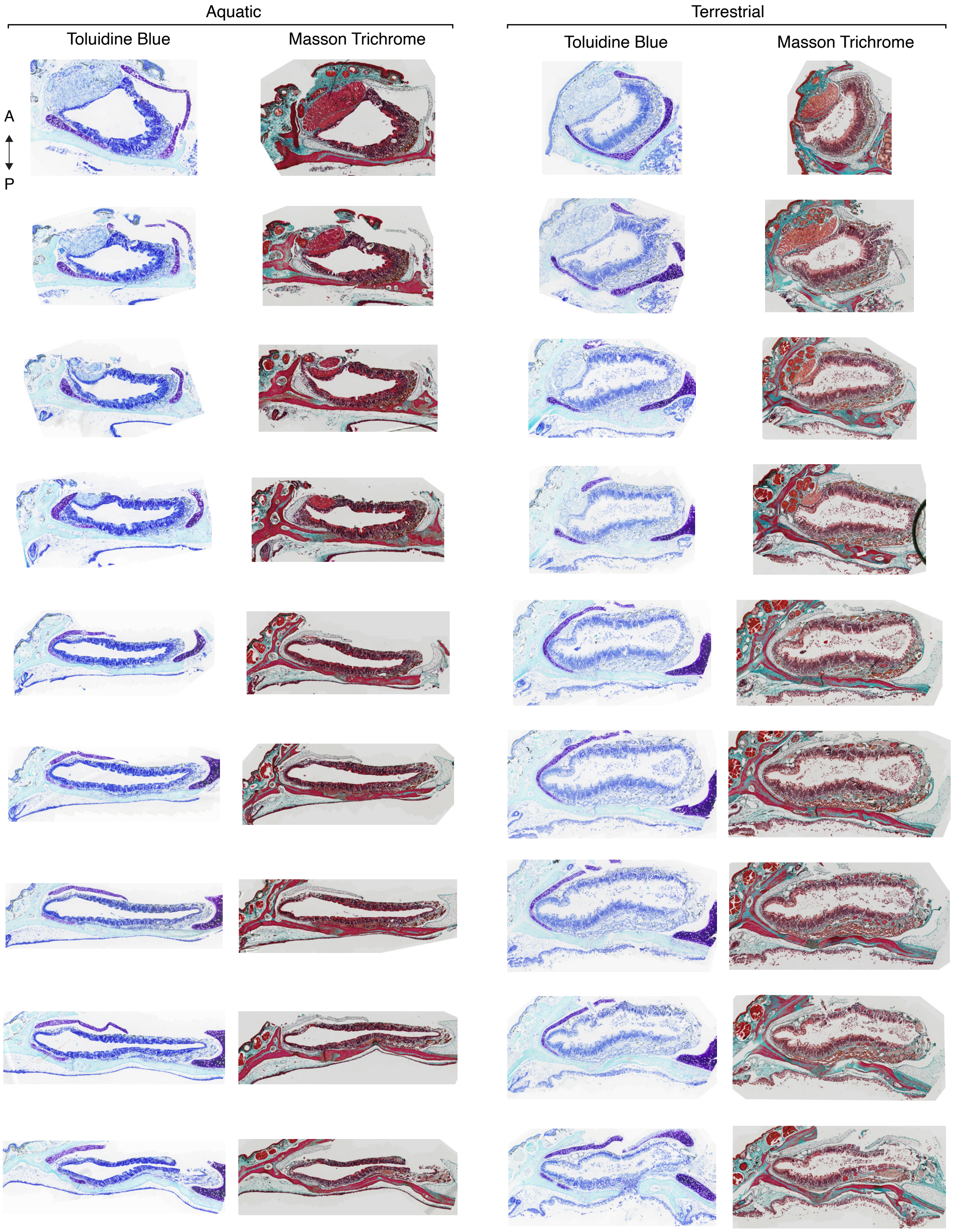

Aquatic
Terrestrial
Toluidine Blue
Masson Trichrome
Toluidine Blue
Masson Trichrome
A
P

**Supplementary Figure 4: Histochemical analysis of the newt nasal epithelium**

A) Serial coronal sections of paraffin-embedded newt olfactory epithelium in aquatic and terrestrial conditions, stained with toulidine blue (left for each condition) or Masson's trichrome (right for each condition). Sections are arranged anterior to posterior.

**A**

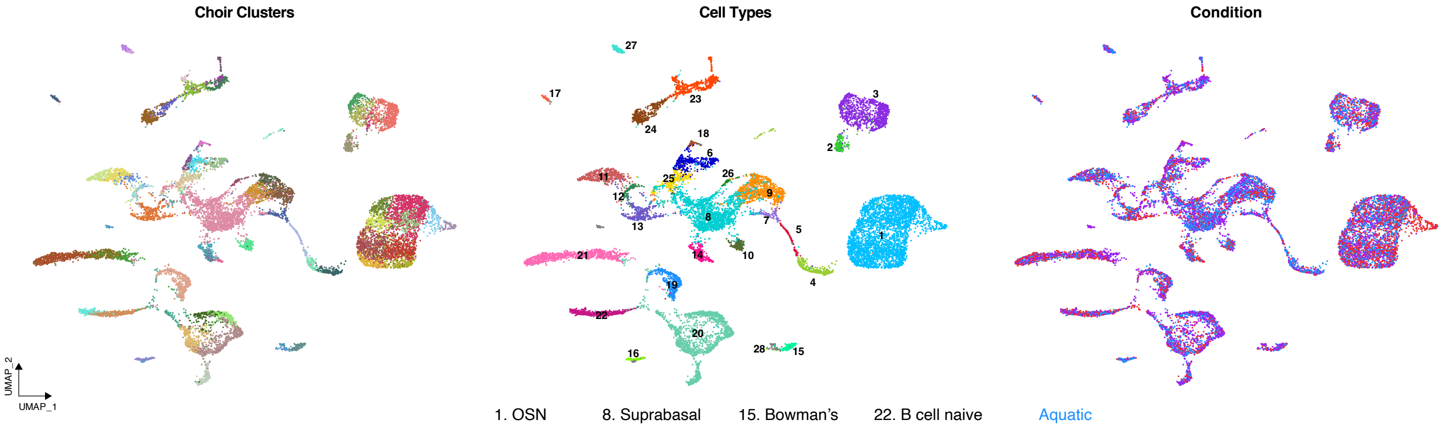


| | | | |
|---|---|---|---|
| 1. OSN | 8. Suprabasal | 15. Bowman's | 22. B cell naive |
| 2. V1R VSN | 9. Sustentacular | 16. Nasal | 23. Fibro |
| 3. V2R VSN | 10. Club | 17. MV Brush | 24. Chondro |
| 4. Imm OSN | 11. Goblet1 | 18. Teeth | 25. Neuromast |
| 5. INP | 12. Goblet2 | 19. Macrophage | 26. Hair Cell |
| 6. HBC | 13. Goblet3 | 20. T cell | 27. HSC |
| 7. GBC | 14. Resp Cil | 21. B cell active | 28. Other |

Aquatic
Terrestrial
Reaquatic

**Supplementary Figure 5: Integration of single-cell RNA sequencing data**
A) (left) UMAP of Integrated nasal epithelium scRNAseq data, clustered using the CHOIR method. (middle) Same UMAP showing cells annotated by type (right) Same UMAP with cells colored by environmental conditions.

A

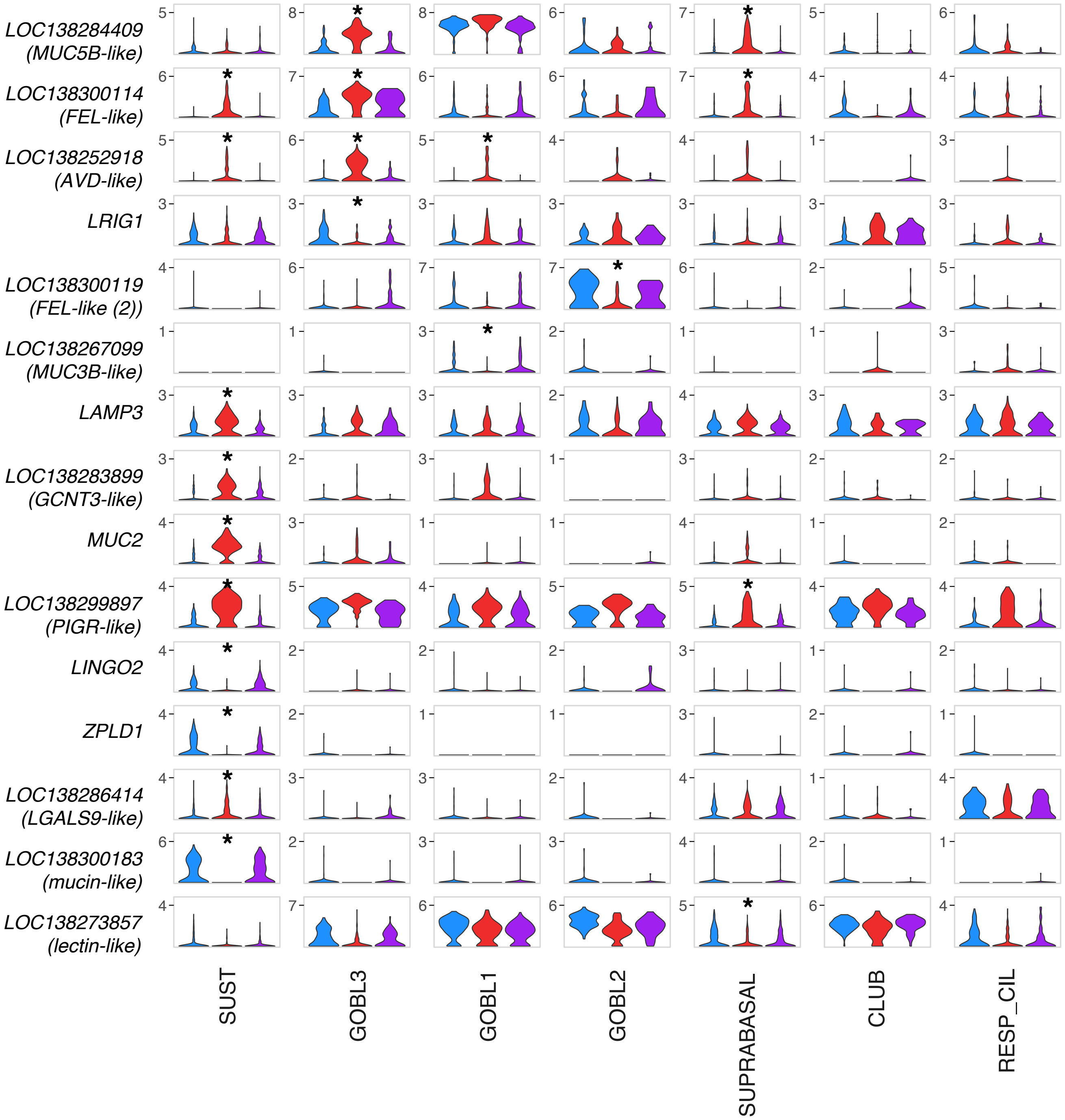
LOC138284409 (MUC5B-like)
LOC138300114 (FEL-like)
LOC138252918 (AVD-like)
LRIG1
LOC138300119 (FEL-like (2))
LOC138267099 (MUC3B-like)
LAMP3
LOC138283899 (GCNT3-like)
MUC2
LOC138299897 (PIGR-like)
LINGO2
ZPLD1
LOC138286414 (LGALS9-like)
LOC138300183 (mucin-like)
LOC138273857 (lectin-like)
SUST
GOBL3
GOBL1
GOBL2
SUPRABASAL
CLUB
RESP_CIL

**Supplementary Figure 6: Mucus related gene expression changes in nasal epithelium non-neuronal cells**

Violin plots showing expression of all mucus related genes (see Methods) with significant differential expression between terrestrial and aquatic conditions in at least one cell type. Star indicates the genes were significantly differentially expressed in that cell type.

A

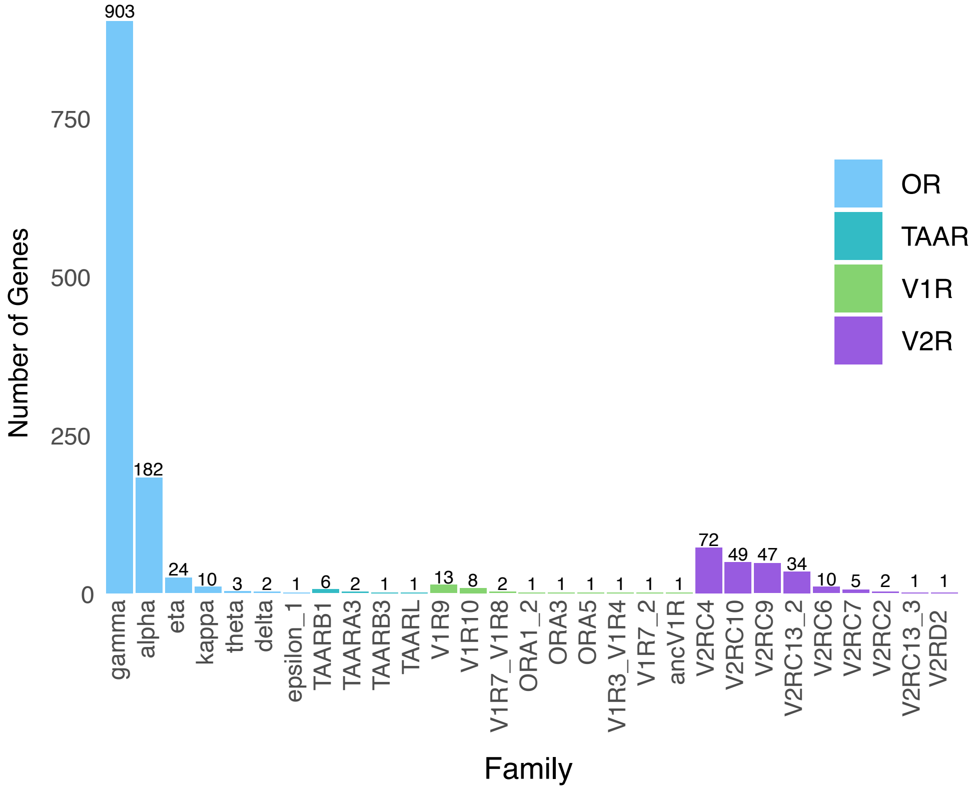

Number of Genes
0
250
500
750
903
182
24
10
3
2
1
6
2
1
1
13
8
2
1
1
1
1
1
1
72
49
47
34
10
5
2
1
1
gamma
alpha
eta
kappa
theta
delta
epsilon_1
TAARB1
TAARA3
TAARB3
TAARL
V1R9
V1R10
V1R7_V1R8
ORA1_2
ORA3
ORA5
V1R3_V1R4
V1R7_2
ancV1R
V2RC4
V2RC10
V2RC9
V2RC13_2
V2RC6
V2RC7
V2RC2
V2RC13_3
V2RD2
Family
OR
TAAR
V1R
V2R

**Supplementary Figure 7: Chemoreceptor gene families in *Pleurodeles waltl***
A) Bar plot showing number of genes identified in the *Pleurodeles waltl* genome for each chemoreceptor gene family using the pipeline described in Policarpo et al., 2024. Numbers above bars indicate the exact number of identified genes from that family.

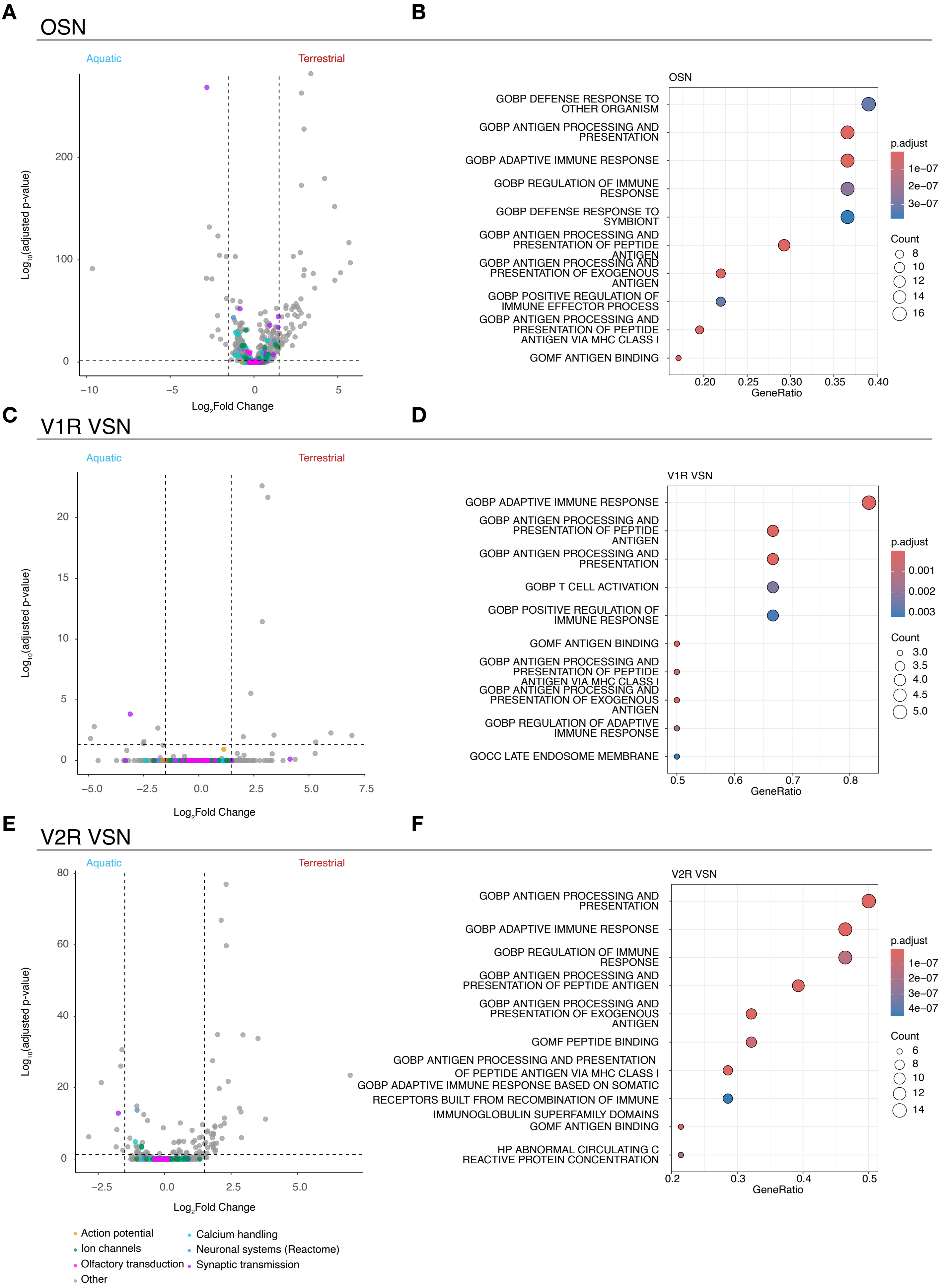
A
OSN
Aquatic
Terrestrial
Log10(adjusted p-value)
Log2Fold Change
B
OSN
GOBP DEFENSE RESPONSE TO OTHER ORGANISM
GOBP ANTIGEN PROCESSING AND PRESENTATION
GOBP ADAPTIVE IMMUNE RESPONSE
GOBP REGULATION OF IMMUNE RESPONSE
GOBP DEFENSE RESPONSE TO SYMBIONT
GOBP ANTIGEN PROCESSING AND PRESENTATION OF PEPTIDE ANTIGEN
GOBP ANTIGEN PROCESSING AND PRESENTATION OF EXOGENOUS ANTIGEN
GOBP POSITIVE REGULATION OF IMMUNE EFFECTOR PROCESS
GOBP ANTIGEN PROCESSING AND PRESENTATION OF PEPTIDE ANTIGEN VIA MHC CLASS I
GOMF ANTIGEN BINDING
GeneRatio
p.adjust
Count
C
V1R VSN
Aquatic
Terrestrial
Log10(adjusted p-value)
Log2Fold Change
D
V1R VSN
GOBP ADAPTIVE IMMUNE RESPONSE
GOBP ANTIGEN PROCESSING AND PRESENTATION OF PEPTIDE ANTIGEN
GOBP ANTIGEN PROCESSING AND PRESENTATION
GOBP T CELL ACTIVATION
GOBP POSITIVE REGULATION OF IMMUNE RESPONSE
GOMF ANTIGEN BINDING
GOBP ANTIGEN PROCESSING AND PRESENTATION OF PEPTIDE ANTIGEN VIA MHC CLASS I
GOBP ANTIGEN PROCESSING AND PRESENTATION OF EXOGENOUS ANTIGEN
GOBP REGULATION OF ADAPTIVE IMMUNE RESPONSE
GOCC LATE ENDOSOME MEMBRANE
GeneRatio
p.adjust
Count
E
V2R VSN
Aquatic
Terrestrial
Log10(adjusted p-value)
Log2Fold Change
F
V2R VSN
GOBP ANTIGEN PROCESSING AND PRESENTATION
GOBP ADAPTIVE IMMUNE RESPONSE
GOBP REGULATION OF IMMUNE RESPONSE
GOBP ANTIGEN PROCESSING AND PRESENTATION OF PEPTIDE ANTIGEN
GOBP ANTIGEN PROCESSING AND PRESENTATION OF EXOGENOUS ANTIGEN
GOMF PEPTIDE BINDING
GOBP ANTIGEN PROCESSING AND PRESENTATION OF PEPTIDE ANTIGEN VIA MHC CLASS I
GOBP ADAPTIVE IMMUNE RESPONSE BASED ON SOMATIC RECEPTORS BUILT FROM RECOMBINATION OF IMMUNE IMMUNOGLOBULIN SUPERFAMILY DOMAINS
GOMF ANTIGEN BINDING
HP ABNORMAL CIRCULATING C REACTIVE PROTEIN CONCENTRATION
GeneRatio
p.adjust
Count
Action potential
Calcium handling
Ion channels
Neuronal systems (Reactome)
Olfactory transduction
Synaptic transmission
Other

**Supplementary Figure 8: The expression of genes related to neuronal function is not altered in sensory neurons sampled from terrestrial and aquatic newts**

A) Volcano plot of log2(Fold Change) and log10(adjusted p-value) for genes expressed in at least 20% of OSNs. Each dot represents a gene, and genes involved in neuronal function, olfactory transduction, and synaptic transmission are highlighted in different colors. Negative fold change values indicate the gene is upregulated in the aquatic condition, and positive values indicate the gene is upregulated in the terrestrial condition.

B) Over representation analysis on differentially expressed genes showing the top 12 gene ontology terms.

C, E) same as A for V2R VSNs and V1R VSNs respectively.

D,F) same as B for V2R VSNs and V1R VSNs.

A

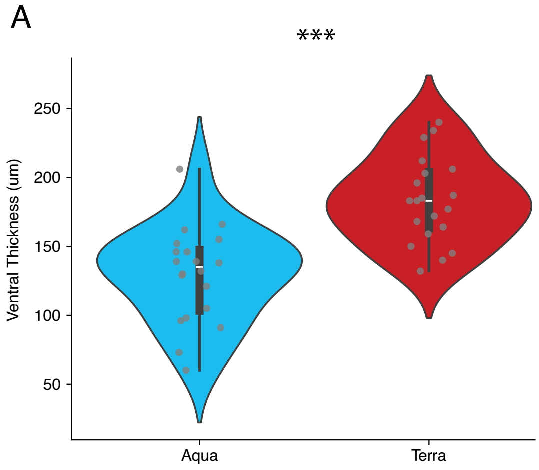
***
Ventral Thickness (um)
250
200
150
100
50
Aqua
Terra

B

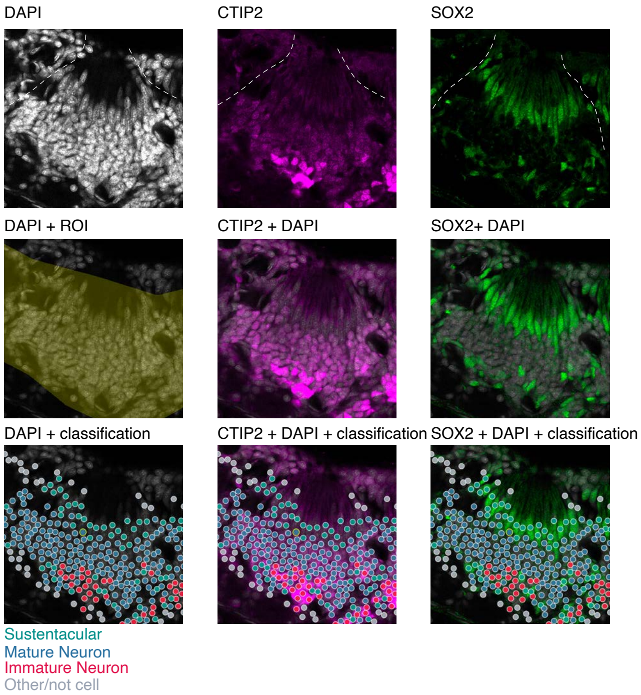
DAPI
CTIP2
SOX2
DAPI + ROI
CTIP2 + DAPI
SOX2+ DAPI
DAPI + classification
CTIP2 + DAPI + classification
SOX2 + DAPI + classification
Sustentacular
Mature Neuron
Immature Neuron
Other/not cell

**Supplementary Figure 9: Quantification of neurogenesis and identification of cell types using Cellpose**

A) Quantification of the thickness of the ventral portion of the olfactory epithelium measurements in terrestrial and aquatic newts. Terrestrial epithelia are significantly thicker than aquatic ones (linear mixed effects model. $p=1.43e-04$).

B) Classification pipeline showing one example olfactory epithelium stripe. (top row) Individual channels showing immunostaining for DAPI (nuclei), CTIP2 (high in immature neurons, moderate in mature neurons) and SOX2 (high in sustentacular cells, GBCs, and INPs). (Middle row) ROI showing region that cell bodies were counted within, and overlays of CTIP2 and SOX2 with DAPI. (bottom row) Overlay with classifications (see materials and methods). Dashed white lines indicate edges of olfactory epithelia stripes.

*Cell-Type abbreviations used in text/figures:*

| Cell Type | Abbreviation | Description |
|---|---|---|
| Olfactory Sensory Neuron | OSN | Olfactory receptor expressing sensory neurons of the nasal epithelium |
| V1 Receptor Type Vomeronasal Sensory Neuron | V1R_VSN | V1 receptor expressing sensory neurons of the nasal epithelium |
| V2 Receptor Type Vomeronasal Sensory Neuron | V2R_VSN | V2 receptor expressing sensory neurons of the nasal epithelium |
| Immature Sensory Neurons | IMM_NEUR | Late development sensory neurons |
| Intermediate Neural Progenitors | INP | Early development sensory neurons |
| Horizontal Basal Cells | HBC | Stem cells of the olfactory and respiratory epithelium |
| Globose Basal Cells | GBC | Progenitors that will adopt an OSN or SUST fate |
| Sustentacular cells | SUST | Epithelial-glia like cells that form the barrier with the external environment with the nose |
| Respiratory ciliated cells | RESP_CIL | Ciliated cells of the respiratory epithelium, sweep particles and mucus through the nasal cavity |
| Suprabasal cells | SUPRABASAL | Immature cells that will develop into one of the support cell lineages |
| Goblet cells | GOBL1/2/3 | Secretory cells that produce mucus and other secretions in the nose |
| Bowman’s glands | BOWMANS | A gland that produces olfactory binding proteins and other secretions |
| Nasal glands | NASAL | Another secretory gland population |

| | | |
|---|---|---|
| Club cells | CLUB | An immature population of secretory cells |
| Microvillous brush cells | MV_BRUSH | Specialized epithelial cells for detecting irritants/foreign compounds |
| Ionocytes | IONO | cells that regulate the fluid balance of the nasal cavity |
| Macrophage | MACRO | Immune cell - defense and clean up |
| T cells | T_CELL | Immune cell |
| B cells (active) | B_CELL_ACTIVE | Immune cell -antigen detection/ antibody production |
| B cells (naive) | B_CELL_NAIVE | Immune cell -antigen detection/ antibody production |
| Fibroblasts | FIBRO | Connective tissue |
| Chondrocytes | CHONDRO | Cartilage cells |
| Dental precursor cells | TEETH | cells that will become (or are) teeth from the palette |
| Neuromast | NEUROMAST | Lateral line cells |
| Hair cells | HAIR_CELL | Cells of the inner ear |
| Hematopoietic stem cells | HSC | immature blood cells |

### *Gene replacements used in text/figures:*

| **Text name** | **Locus identifier** | **Description** |
|---|---|---|
| MUC5B-like | LOC148284409 | mucin-5B-like |
| RSN-like | LOC138260829 | ranaspumin-like |
| PIGR-like | LOC138299897 | Polymeric immunoglobulin receptor-like |
| LCN15-like | LOC138302175 | lipocalin-15-like |
| HMGN5-like | LOC138247144 | high mobility group nucleosome-binding domain-containing protein 5-like |

| | | |
|---|---|---|
| CD72-like | LOC138296275 | B-cell differentiation antigen CD72-like |
| CYP3A4-like | LOC138261214 | cytochrome P450 3A21-like (non-human primate equivalent to CYP3A4) |
| MUC5AC-like | LOC138284405 | mucin-5AC-like |
| AVD-like | LOC138252918 | avidin-like |
| SULT1B1-like | LOC138304187 | sulfotransferase 1B1-like |
| FEL-like | LOC138300114 | fish-egg lectin-like |
| FEL-like (2) | LOC138300119 | fish-egg lectin-like |
| CFB-like | LOC138299143 | complement factor B-like |
| CAT-like | LOC138283776 | catalase-like |
| FTH1-like | LOC138246336 | ferritin heavy chain B |
| CASP7-like | LOC138300243 | caspase-7-like |
| CASP7-like (2) | LOC138300242 | caspase-7-like |
| PARP14-like | LOC138284950 | protein mono-ADP-ribosyltransferase PARP14-like |
| IFI44L-like | LOC138293160 | interferon-induced protein 44-like |
| CXCL10-like | LOC138298109 | C-X-C motif chemokine 10-like |
| MUC3B-like | LOC138267099 | Mucin-3B-like |
| GCNT3-like | LOC138283899 | beta-1,3-galactosyl-O-glycosyl-glycoprotein beta-1,6-N-acetylglucosaminyltransferase 3-like |
| LGALS9-like | LOC138286414 | galectin-9-like |
| Lectin-like | LOC138273857 | lectin-like |
